# Stimulation with mycobacterial glycolipids and PPD reveals different innate immune response profiles in active and latent TB

**DOI:** 10.1101/2021.04.04.434373

**Authors:** Carolina S Silva, Christopher Sundling, Elin Folkeson, Gabrielle Fröberg, Claudia Nobrega, João Canto-Gomes, Benedict J. Chambers, Tadepally Lakshmikanth, Petter Brodin, Judith Bruchfeld, Jerome Nigou, Margarida Correia-Neves, Gunilla Källenius

## Abstract

Upon infection with *Mycobacterium tuberculosis* (Mtb) the host immune response might clear the bacteria, control its growth leading to latent tuberculosis (LTB), or fail to control its growth resulting in active TB (ATB). There is however no clear understanding of the features underlying a more or less effective response. Mtb glycolipids are abundant in the bacterial cell envelope and modulate the immune response to Mtb, but the patterns of response to glycolipids are still underexplored. To identify the CD45^+^ leukocyte activation landscape induced by Mtb glycolipids in peripheral blood of ATB and LTB, we performed a detailed assessment of the immune response of PBMCs to the Mtb glycolipids lipoarabinomannan (LAM) and its biosynthetic precursor phosphatidyl-inositol mannoside (PIM), and PPD. At 24 h and 5 days of stimulation, cell profiling and secretome analysis was done using mass cytometry and high-multiplex immunoassay. PIM mainly affected antigen-presenting cells to produce both proinflammatory (IL-2, IL-6, IL-17A, TNF-*α* and GM-CSF), and IL-4 and IL-10 cytokines, but not IFN-γ. LAM triggered a similar, albeit weaker, response. By contrast, PPD induced an increase in IFN-γ-producing cells. Moreover, PPD also led to increased numbers of IL-2, IL-6, IL-10, IL-17A, TNF-*α* and GrzB-producing cells. Treatment with an anti-TLR2 antibody led to partial inhibition of PIM-induced IL-6 production in myeloid cells, suggesting that PIM induces IL-6 production through TLR2. Expansion of monocyte subsets in response to PIM or LAM was reduced in both ATB and LTB as compared to healthy controls, suggesting a hyporesponsive/tolerance pattern in Mtb-infected individuals.

## Introduction

Approximately 10 million people develop tuberculosis (TB) each year, and it is estimated that around 25% of the world population is latently infected with *Mycobacterium tuberculosis* (Mtb) (1). However, only about 10% of individuals with latent TB (LTB) are estimated to develop active TB (ATB) (2). It is clear therefore that in most cases Mtb infection is well controlled, but our understanding of what makes an effective immune response that controls and/or clears Mtb is limited.

Research on the host response to Mtb has so far mainly focused on antigens of protein/peptide nature. However, the immune response to Mtb is initiated mainly through the interaction of Mtb cell envelope components, mostly glycolipids, with cells of the innate immune system (3), which trigger activating or repressive responses in terms of cytokine production (4, 5). Glycolipids are abundantly expressed in all mycobacterial species. The ability of Mtb lipids to traffic outside infected cells (6-8) renders the direct contact of Mtb cell envelope glycolipids with distinct immune cells an important aspect of the immune response (9). Lipoarabinomannan (LAM) is a major glycolipid of the Mtb cell wall and has been studied quite extensively for its immunomodulatory properties (10, 11), compared to its biosynthetic precursors, the phosphatidyl-inositol mannosides (PIM_2_, and PIM_6_). A large number of host cell receptors take part in the initial interaction between mycobacteria and innate immune cells (12, 13). TLRs and C-type lectins are involved in this process, resulting in activation of several antimicrobial mechanisms by macrophages (Mφs) and dendritic cells (DCs) (14-17).

In addition to the extensive interaction with innate immune cells, PIM and LAM are also both recognized by CD1b-restricted T cells (9, 18-21). In fact, it was observed that purified-protein derivate (PPD) positive individuals respond through CD1-restricted T cells to several mycobacterial lipids, including PIM and LAM (18) and that this response may vary between individuals with ATB and LTB. Mtb whole lipid extract was shown to induce proliferation of CD1-restricted CD4^+^ and, to a smaller extent, CD8^+^ T cells in LTB. Interestingly, the same was observed for ATB patients only after the first two weeks of anti-TB treatment (22). A subset of LAM reactive CD1-restricted T cells co-expressing perforin, granulysin, and granzyme B (GrzB), mostly CD8^+^, are more frequent in LTB than in individuals who developed active disease after infection (evaluated after TB treatment) (22). Similarly, glycerol monomycolate-specific T cells are more frequent in LTB than ATB patients (18) and the response of these cells may vary between ATB and LTB individuals.

B cell-mediated immunity in Mtb infection has been less explored compared to monocyte- and T cell-mediated responses, although recent data strengthen the relevance of these cells in the immune response to Mtb. Recently it was shown that Mtb LAM induces IL-10 production by B cells and that these cells (B10) inhibit CD4_+_ T_H_1 polarization leading to increased Mtb susceptibility in mice (23). The response of B cells to LAM was shown to occur in a TLR2-dependent manner (23). Stimulation with LAM has also been reported to lead to IL-4 production by CD19^+^/B220^+^ precursor cells in the bone marrow of mice, presumably pre-B cells (24).

In the present study, we performed a detailed assessment and simultaneous comparison of the immune response to PIM, LAM and PPD from Mtb in peripheral blood mononuclear cells (PBMCs) from individuals with active or latent TB and compared with healthy controls (HC). We performed immune profiling by secretome analysis and mass cytometry measuring simultaneously 37 cellular markers at the single-cell level to allow high-resolution of the cellular composition and secretion. We identified distinct subsets within memory T cells, NK cells, B cells and monocytes/DCs that were altered by PPD, PIM and LAM stimulation and further evaluated the role of TLR2 in this process.

## Materials and Methods

### Study participants

Participants were recruited in 2018 within an ongoing prospective cohort of adult (≥18 years) TB patients and contacts attending the TB Centre, Dept of Infectious Diseases Karolinska University Hospital Stockholm (Supplemental Table A). Active TB (ATB) cases were defined upon microbiological (PCR and/or culture) verification. Latent TB (LTB) participants were defined as asymptomatic, IGRA positive, close contacts to ATB cases. Healthy controls (HC) were defined as IGRA negative students and hospital staff without known previous TB exposure. Exclusion criteria were pregnancy, autoimmune diseases and HIV co-infection or other immunodeficiencies. ATB and LTB participants were screened with standard biochemical set-up and radiology.

### Antigens

Tuberculin PPD (RT 50) was obtained from Statens Serum Institute, and PHA from Invivogen. LAM and PIM were prepared in-house (5). PIM contains both PIM_2_ and PIM_6_ isoforms, differing in number of fatty acyl constituents (5).

### PBMC isolation

Venous blood from each participant was collected into EDTA tubes and peripheral blood mononuclear cells (PBMCs) were purified through density gradient centrifugation using Lymphoprep™ (Stemcell) according to the manufacturer’s instruction, with some modifications. Briefly, white blood cells were counted using a HemoCue instrument and the blood was diluted to a maximum of 240×10^6^ cells per 22.5 ml that were then layered onto 10 ml Lymphoprep. The cells were centrifuged at 400*g* for 30 min without any break. The mononuclear cell layer was collected into a new 50 ml tube and resuspended to 45 ml with PBS. The cells were spun at 300g for 10 min with break after which the cells were resuspended into 1-5 ml PBS and filtered using a 100 μm poor size cell strainer and counted on a Countess (ThermoFisher Scientific). The cells were centrifuged at 400*g* for 10 min with break and resuspended with freeze media (90% FBS supplemented with 10% DMSO) and placed in a CoolCell freezing container (Sigma) before moving to –80°C overnight followed by long-term storage at liquid nitrogen.

### PBMCs stimulation

PBMCs from five patients with ATB, five with LTB and five HCs were thawed at 37°C followed by addition of 1 mL RPMI-1640 media supplemented with 10% fetal bovine serum (FBS), 1% penicillin/streptomycin (P/S) and 250 U/mL Benzonase (all from ThermoFisher). The cells were washed twice (300*g* for 5 min) in media followed by resuspension in RPMI-1640 culture media supplemented with 10% FBS, 1% P/S, 0.3 g/L L-Glutamine and 25 mM HEPES and counted. The cells were then plated in 24-well plates at 2×10^6^ PBMCs/mL in culture media containing either 5 μg/mL PHA, 10 μg/mL PPD, or 25 μg/mL LAM or PIM, or left untreated (PBS), for 24 hours or 5 days in a 37°C 5% CO_2_ incubator. Four hours before collection, 5 μg/mL of brefeldin A and 2 μM Monensin (both ThermoFisher) were added to each well. PHA was used as positive control for PBMCs responsiveness (Supplemental Figure 1 and 2). The choice of concentration of LAM and PIM was based on previous unpublished work.

### Mass cytometry staining and acquisition

After 24 h and 5 days of stimulation, cells were collected by centrifugation after a 15 min incubation with 2 mM EDTA. Supernatants were stored at –80°C and cells were fixed using the PBMCs fix kit (Cytodelics AB) and barcoded using Cell-ID™ 20-Plex Pd Barcoding Kit (Fluidigm Inc.), according to the manufacturer’s recommendations. Samples were washed with CyFACS buffer (PBS with 0.1% BSA, 0.05% sodium azide and 2mM EDTA) and Fc receptors were blocked with 200 μL of blocking buffer (Cytodelics AB) for 10 min at RT. Cells were incubated with 200 μL of antibody cocktail (Supplemental Table 2) for 30 min at 4°C, washed with CyFACS buffer, and fixed with 1% formaldehyde. For intracellular staining, cells were permeabilized using an intracellular fixation and permeabilization kit (eBiosciences Inc.) according to the manufacturer’s instructions. Subsequently, 200μl of intracellular antibody cocktail (Supplemental Table 3) was added and incubated for 45 min at RT. Cells were washed, fixed in 4% formaldehyde at 4°C overnight, and stained with DNA intercalator (0.125 µM MaxPar® Intercalator-Ir, Fluidigm Inc.) on the following day. After that, cells were washed with CyFACS buffer, PBS and MiliQ water, counted and adjusted to 750,000 cells/mL. Samples were acquired in a CyTOF2 (Fluidigm) mass cytometer at a rate of 250-400 events/s using CyTOF software version 6.0.626 with noise reduction, a lower convolution threshold of 200, event length limits of 10-150 pushes, a sigma value of 3, and flow rate of 0.045 ml/min.

### Analysis of mass cytometry data

The mass cytometry FCS data files were gated for different cell subsets: CD45^+^ leucocytes, CD45^+^CD3^+^CD20^−^ T cells, CD45^+^CD3^−^CD7^+^ NK cells, CD45^+^CD3^−^HLA-DR^+^ antigen-presenting cells (APCs), and CD45^+^ leukocytes producing IL-2, IL-4, IL-5, IL-6, IL-10, IL-17A, IFN-γ, TNFa, GrzB, and GM-CSF using FlowJo™ v10.6.1. The gated populations were exported to new FCS files that were then analysed using the R-package Cytofkit v1.12.0, which includes an integrated pipeline for mass cytometry analysis (25). Cytofkit was run in R-studio version 1.1.463 and R version 3.6.1. For analysis of total leukocytes, 5000 cells were used per sample. For analysis of gated T cells, NK cells, and APCs, 10000 cells were used per sample. For analysis of cytokine^+^ cells, a ceiling of 5000 cells were included per sample. Dimensionality was reduced using Barnes-Hut tSNE with a perplexity of 30 with a maximum of 1000 iterations. Clustering was then performed using density-based machine learning with ClusterX (25) and cell subsets were identified by visual inspection of marker expression for each cluster. The Cytoftkit analysis was performed using PBS, PPD, PIM, and LAM FCS files together, whereas PHA stimulated cells were evaluated independently, using only PBS and PHA FCS files.

### Secretome analysis of culture supernatants

Cell culture supernatants (n=75) were randomized in a 96-well plate and analyzed with a multiplex proximity extension assay (PEA) (26), enabling simultaneous quantification of 92 inflammatory markers from the Olink inflammation panel (Supplemental Table 4). Markers where all samples were below the limit of detection of the assay were removed from subsequent analysis. The samples were run by the Translational Plasma Profile Facility at SciLifeLab, Stockholm, Sweden.

### TLR2-dependence of PBMCs activation

To investigate TLR2-dependent PBMCs activation by the Mtb glycolipids PIM and LAM, frozen PBMCs from HC (n=5) were thawed in a 37°C water bath, washed two times in complete media (RPMI-1640 culture media supplemented with 10% FBS, 1% P/S, 1mM sodium pyruvate and 10 mM HEPES) and plated as described for mass cytometry. Prior to stimulation, the cells were pre-incubated for 30 min at 37°C with 5 μg/mL of anti-TLR2 monoclonal antibody (clone T2.5, InvivoGen) or with an isotype control (mIgG1, eBiosciences).

### Flow cytometry

Cells stimulated in the presence or absence of anti-TLR2 antibody for 24 h were collected after an additional 15 min incubation with 2mM EDTA. The cells were then washed with FACS buffer (PBS with 0.3% BSA and 2mM EDTA) and Fc receptors were blocked with 20 μL of blocking buffer Fc Receptor Binding Inhibitor (eBiosciences) for 10 min at 4 °C. The cells were incubated with 50 μL of antibody cocktail (Supplemental Table 5) for 30 min at RT, washed with PBS and incubated with Fixable Viability Dye eFluor™ 450 (eBiosciences) for 30 min at 4 °C. For intracellular staining, the cells were permeabilized using the FoxP3 intracellular fixation and permeabilization kit (eBiosciences) according to the manufacturer’s instructions. Subsequently, 50 μl of intracellular Ab cocktail (Supplemental Table 5) was added and incubated for 30 min at 4 °C. Finally, the cells were washed, resuspended in PBS and kept at 4°C until acquisition on the next day. The cells were acquired on a 12-color LSRII flow cytometer using FACSDiva software (Becton Dickinson, Franklin Lakes, NJ); data analysis was performed using FlowJo™ v10.6.1. Gating strategies are represented in Supplemental Figure 3.

### Statistical analysis

Comparisons of a single variable for paired data for >2 groups were evaluated by Friedman’s test followed by Dunnet’s post-hoc test. Comparisons of a single variable for unpaired data for >2 groups were evaluated by using a Kruskal-Wallis test followed by Dunn’s post-test. Comparisons of >1 variable for paired data were evaluated using repeated measures 2-way ANOVA followed by Dunnett’s post-hoc test. Differences were considered significant when p<0.05. Statistical analyses were performed using Prism9 (GraphPad Software, USA).

### Study approval

Written informed consent was received from all participants before inclusion in the study. The study was approved by the Regional Ethical Review Board at the Karolinska Institute in Stockholm (approval numbers 2013/1347-31/2 and 2013/2243-31/4) and by the Ethics Committee for Research in Life and Health Sciences of the University of Minho, Portugal (approval number SECVS 014/2015) and it is in accordance with the Declaration of Helsinki.

## Results

### Effect of stimulants on cell types and cytokine production

To investigate the effect of the Mtb glycolipids LAM and PIM on the immune response, PBMCs from individuals with ATB or LTB, and HC (Supplemental table 1) were stimulated for 24 h and 5 days; PPD was used as a control for responses to Mtb proteins, while mock stimulation (PBS) or phytohemagglutinin (PHA) were used as negative and positive culture controls, respectively. Proteins released into the culture supernatant were analysed using the Olink proximity-extension assay (PEA), that allows for simultaneous measurement of 92 inflammatory markers. Cells were analysed using mass cytometry for changes in the expression of 27 surface and 10 intracellular markers (Figure 1 and Supplemental table 2 and 3).

**Figure 1.**
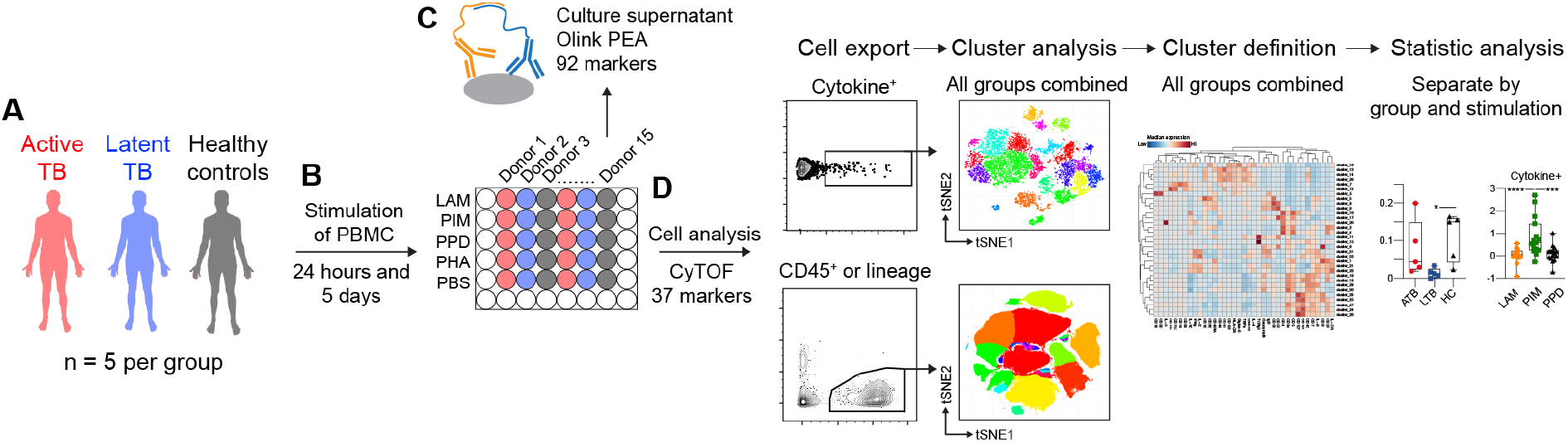
Schematic of the study outline. (**A**) Three groups, including active TB (ATB, red), latent TB (LTB, blue) and healthy controls (HC, black) were (**B**) stimulated for 24 hours or 5 days with the the mycobacterial glycolipids lipoarabinomannan (LAM) or phosphatidyl-inositol mannoside (PIM). Control stimulations included purified protein derivate (PPD), or phytohemagglutinin (PHA), or mock stimulation (PBS). (**C**) Culture supernatants were analysed by proximity extension assay (PEA), while (**D**) cells were analysed by mass cytometry. Cells were either pre-gated for cytokine-secreting cells or different cell subsets prior to dimensionality reduction and cluster analysis.

Cytokines are rapidly produced from both innate and adaptive immune cells following stimulation with bacterial components. The pattern of cytokines that are produced indicates the overall polarization of the immune response, which can affect pathogen control (27). To assess the early (24 hours) and late (5 days) effect of LAM, PIM, and PPD on secretion of cytokines and chemokines from stimulated PBMCs, we assessed the culture supernatant for relative levels of 92 different soluble inflammatory markers (Figure 2). Of these, we observed changes in protein levels for 60 proteins at 24 hours and for 71 proteins at 5 days of culture. LAM and PIM stimulation produced very similar marker profiles at both 24 hours and 5 days of culture, respectively, with a slightly stronger effect from PIM, suggesting a similar mechanism of action. PPD induced a markedly different response, primarily seen at 5 days of culture, although IFN-γ levels were already considerably higher at 24 hours compared to LAM and PIM, potentially suggesting a different mechanism of action. At 24 hours there were some indications of different levels of secretion between the groups (ATB/LTB/HC), primarily with a greater effect observed for HC compared with ATB or LTB (Figure 2A). This effect then became more pronounced at 5 days of stimulation (Figure 2B), especially for PPD, where effects in the supernatants from individuals with ATB and/or LTB were significantly higher for CXCL9, CXCL10, CXCL11, IL-13, IL-17A, LIF, IFN-γ, TNF-β, CSF-1, and HGF compared to HC (Figure 2B). Stimulation with LAM and PIM displayed a contrasting pattern, where individuals with LTB and ATB had a less pronounced response compared to HC. At 24 hours IL-1α, IL-10, IL-18, and VEGF were detected at higher levels in HC compared with LTB or ATB, while no such increase was observed between the groups at 5 days. Instead, in several instances, LAM and/or PIM stimulation led to lower protein levels than what was observed for unstimulated cells (dotted line), potentially indicating that the proteins were consumed over time or that production was blocked. Interestingly, for several proteins (CCL8 at 24 hours and CCL13, CXCL9, 10, 11, IL-17A, uPA, and TWEAK at 5 days) this effect was more pronounced in HC compared with ATB and/or LTB. Several of the proteins with negative responses function as chemoattractants (especially for monocytes), suggesting that their lower levels could be due to their increased uptake from the culture supernatant by activated monocytes.

**Figure 2.**
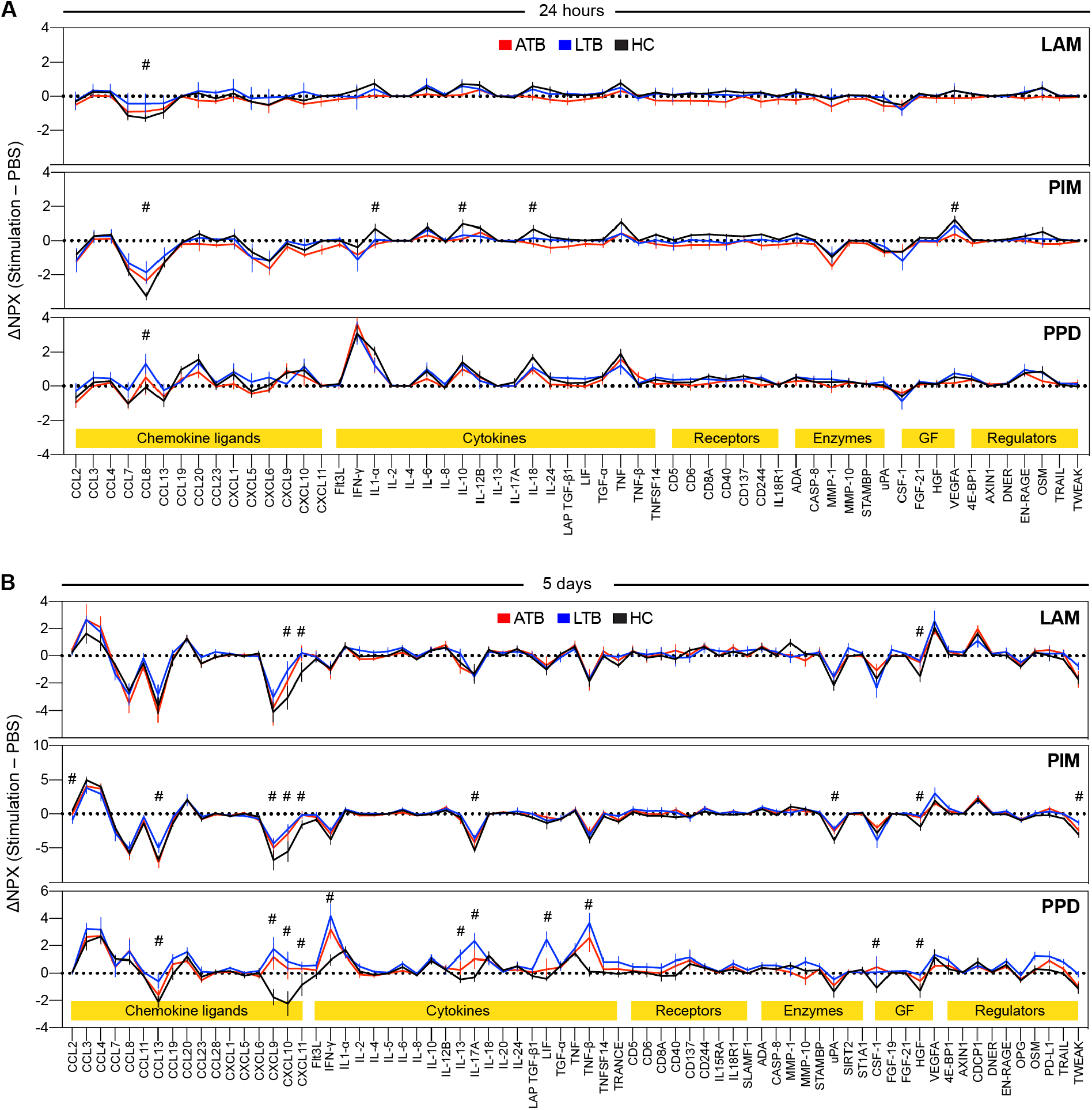
Secretion of inflammatory proteins after stimulation with LAM, PIM, and PPD. Culture supernatants were assessed for 92 protein markers using the Olink inflammation panel after (**A**) 24 hours and (**B**) 5 days of culture. Only markers above background levels were included (n=60 at 24 hours and n=71 at 5 days). The graphs show normalized protein expression (ΔNPX) for stimulated (LAM, PIM, or PPD) minus unstimulated (PBS) PBMCs from individuals with active TB (ATB; red), latent TB (LTB, blue), and healthy controls (HC, black). The lines indicate the mean and error bars indicate SEM. The protein markers were grouped into different functional groups (yellow highlight), where GF correspond to growth factors. Statistics was evaluated using two-way ANOVA followed by Dunnet’s posthoc test with significant differences between ATB or LTB against HC indicated by #, corresponding to a p-value < 0.05.

As the different stimulations could affect cell survival, we assessed the total number of leukocytes upon stimulation, identified through the expression of CD45 (Supplemental Figure 4). After 24 h of stimulation, the number of leukocytes was not changed for LAM, PIM, or PPD compared with unstimulated cells. After 5 days, however, the number of CD45^+^ cells was reduced in stimulations with LAM and in a more pronounced fashion with PIM.

### Intracellular cytokine production in response to stimulation

To evaluate the effect of each stimulus on intracellular production of cytokines and GrzB, regardless of the experimental group, the cumulative frequency of cytokine^+^ cells among CD45^+^ leukocytes was compared to that of unstimulated cells (PBS; Figure 3A and 3B). As expected, PPD stimulation resulted in an increase in IFN-γ-producing cells both at 24 h and 5 days of stimulation. Moreover, it also led to higher levels of IL-2, IL-6, IL-10, IL-17A, TNF-*α* and GrzB-producing cells. Both IL-6^+^ and IL-10^+^ cells remained higher at 5 days of stimulation, while there was also an increase in GM-CSF-producing cells. Unlike PPD, PIM did not stimulate production of IFN-γ but instead stimulated early production of IL-4 and GM-CSF. In addition, 24 h PIM stimulation led to increased levels of IL-2^+^, IL-6^+^, IL-10^+^, IL17A^+^and TNF-*α*^+^ cells (Figure 3B). At 5 days, IL-10^+^ cells remained elevated while IL-2^+^, TNF-*α*^+^, and GrzB^+^ cells were decreased by PIM stimulation in comparison to unstimulated cells. LAM stimulation resulted in reduced levels of IFN-γ ^+^, IL-4^+^, IL-5^+^, IL-17A^+^, TNF-*α*^+^, and GrzB^+^ cells at 5 days of stimulation. This reduction in cytokine-producing cells in LAM and PIM stimulated cells is potentially due to the strong contraction of live cells after 5 days of stimulation (Supplemental Figure 4A). To get an overview of which cell types that were responsible for the cytokine production, we identified the cell subsets producing each cytokine, regardless of the group and stimuli. We observed that myeloid cells (identified through the expression of CD33) contributed strongly to the early production of IL-2, IL-6, IL-10, TNF-*α*, and GM-CSF. This is consistent with myeloid cells being the main source of pro-inflammatory cytokines such as IL-6 and TNF*α* (28) (Figure 3C). This pattern largely overlapped with the cytokines stimulated by PIM, indicating that myeloid cells could be the main effector cells stimulated by Mtb glycolipids. T cells were the main cytokine producers of IFN-γ, IL-4, IL-5, IL-17A, and GrzB at both 24 h and 5 days. They also took over from myeloid cells for IL-6 and IL-10 at 5 days (Figure 3C). In contrast, NK cells primarily produced cytokines at the early time-point, with a strong contribution to IFN-γ, IL-2, IL-6, IL-17A, TNF*α*, and GrzB-producing cells. Of these at 5 days, NK cells retained a similar proportion only of IL-6 and GrzB-secreting cells. We also identified B cells, producing primarily IL-2, IL-4, IL-5, IL-17A, and TNF*α*, although to a smaller extent compared with the other cell subsets (Figure 3C).

**Figure 3.**
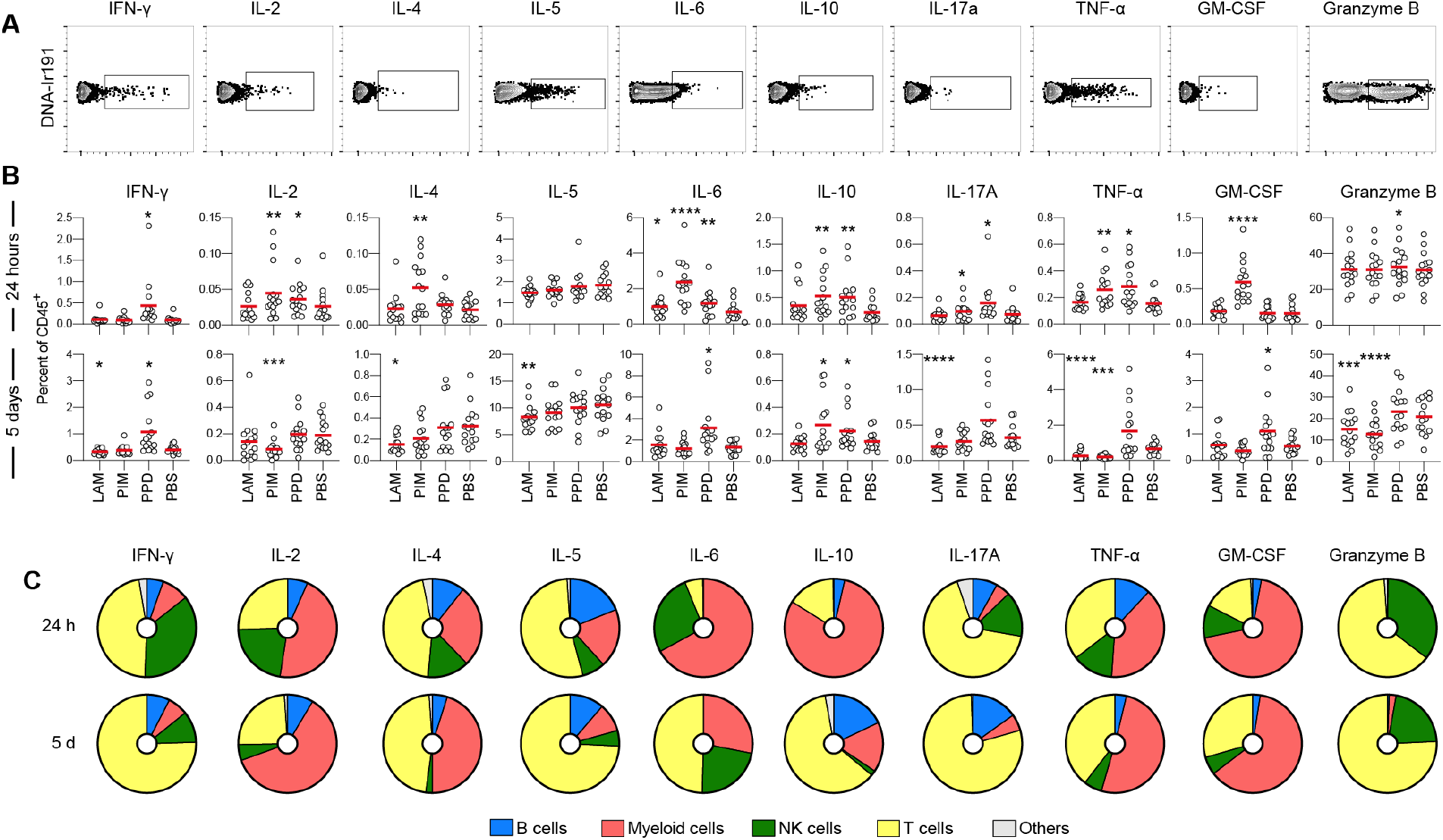
Stimulation-induced cytokine production. (**A**) Representative gating strategy for identification of cytokine-producing CD45^+^ cells via intracellular staining. (**B**) The frequency of cytokine positive cells out of CD45+ cells for each stimulation at 24 hours (top) and 5 days (bottom). All donors merged (n=15/stimulation). Statistics was evaluated by Friedman’s test with Dunnet’s posttest where every group was compared with unstimulated cells (PBS). *p<0.05, **p<0.01, ***p<0.001, ****p<0.0001. (**C**) For each cytokine, the pie-charts indicate the frequency of cell type responsible for its production: B cells (blue), myeloid cells (red), NK cells (green), T cells (yellow), and others (grey).

In summary, stimulation with the Mtb glycolipids LAM and PIM led to early production of several cytokines in multiple cell subsets. The contribution of the individual cell subsets to each cytokine was dynamic over time, potentially indicating inherent differences in cell activation. However, the reduction in different cell subsets over time after stimulation complicates interpretations of the day 5 time-point data. We, therefore decided to focus on understanding the impact of glycolipid stimulation on the early 24 h time-point, with a few exceptions, when overall cell numbers remained unaffected by the stimulation.

### Reduced cytokine production in individuals with active- or latent TB

To investigate the overall cytokine response profile of the main cell populations in individuals with active or latent TB and HC, we pooled all the cytokine-producing cells of each cell population and compared the cumulative production of cytokines within the different groups (Figure 4). For T cells, we observed a reduced cytokine production in individuals with ATB and LTB to PIM stimulation, mainly due to reduced production of GrzB (Figure 4A). There was no overall significant effect on cytokine^+^ NK cells associated with Mtb-infection (Figure 4B). For B cells, the overall cytokine production was reduced in individuals with ATB upon LAM and PIM stimulation compared with HC, primarily due to a reduced production of IL-5 (Figure 4C). For myeloid cells, a similar reduction of cytokine^+^ cells was observed in individuals with LTB to LAM, PIM, and PPD stimulations. This effect was mainly attributed to a reduced production of IL-10 and IL-6. For ATB this effect was only observed in response to LAM (Figure 4D).

**Figure 4.**
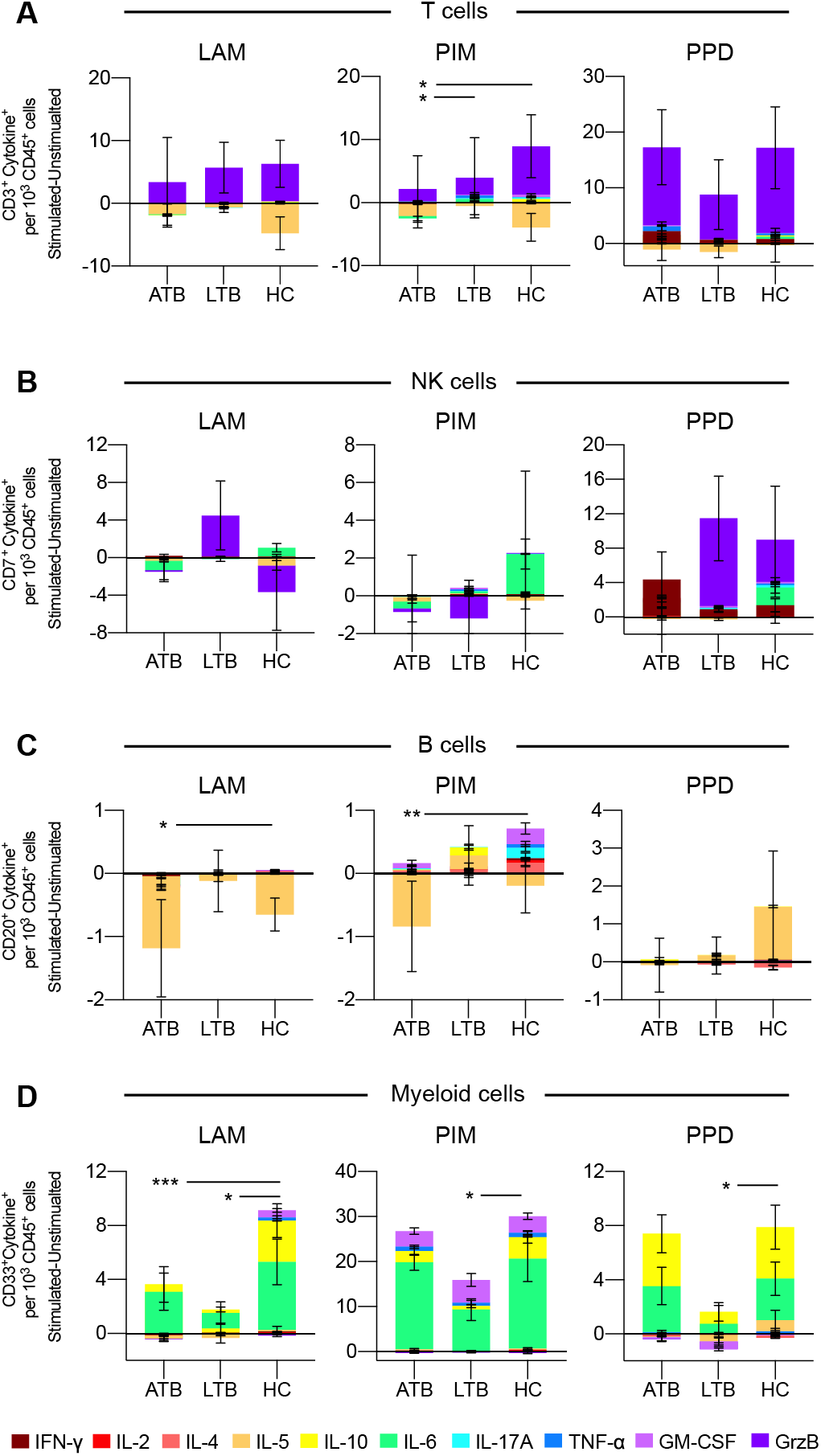
Overall effect of stimulation on cytokine production from individuals with active- and latent TB and healthy controls. Average number of cytokine^+^ cells following 24 h of stimulation with LAM, PIM, or PPD of PBMCs from individuals with active TB (ATB), latent TB (LTB), or healthy controls among (**A**) CD3^+^ T cells, (**B**) CD3^−^CD7^+^ NK cells, (**C**) CD20^+^ B cells, and (**D**) CD33^+^ myeloid cells. The groups average number of cytokine-producing cells were compared using the Friedman test followed by Dunn’s posttest. *p<0.05, **p<0.01, ***p<0.001, ****p<0.0001

In summary, individuals with ATB or LTB responded with less cytokine production by especially myeloid cells and somewhat by B and T cells upon stimulation with Mtb antigens.

To further analyse the effect of LAM, PIM, and PPD on cytokine production by individual cell subsets between the three groups (ATB, LTB, and HC), we proceeded with dimensionality reduction using t-stochastic neighbour embedding (t-SNE) and cluster analysis. This was performed by pooling all PBS, PPD, PIM, and LAM mass cytometry data files together followed by analysis using Cytofkit (25). To allow a high level of resolution in the analysis, cytokine-producing CD45^+^ cells were gated for the individual cytokines (see gates in Figure 3A) which were then analysed separately (Supplemental Figure 5 and 6).

### Qualitatively different T cell responses to PIM and PPD

Stimulation with PPD resulted in an increased number of T cells (identified as CD3^+^) producing IFN-γ, IL-2, IL-5, IL-6, IL-17A, TNF-*α*, and GrzB compared with LAM and/or PIM. Stimulation with PIM contributed to higher numbers of IL-2^+^, TNF-*α*^+^, and GM-CSF^+^ T cells compared with LAM stimulated cells (Figure 5A). Of all cells producing IFN-γ at 24 h, T cells represented 39%, comprising 11 different clusters (clusters 2, 3, 5, 8, 10, 11, 12, 14 15, 16, and 24) (Figure 5B). Four of these clusters (clusters 5, 11, 12, and 15) were significantly elevated by PPD stimulation compared to PIM and/or LAM (Figure 5C). These clusters corresponded to different CD4^+^ and CD8^+^ T cells subsets, including central memory (CD4^+^CD45RA^−^CD27^+^, cluster 5), effector memory (CD45RA^−^CD27^−^, clusters 11 and 15), and effector memory T cells re-expressing CD45RA (TEMRA - CD8^+^CD45RA^+^CD27^−^, cluster 12; Figure 5D).

**Figure 5.**
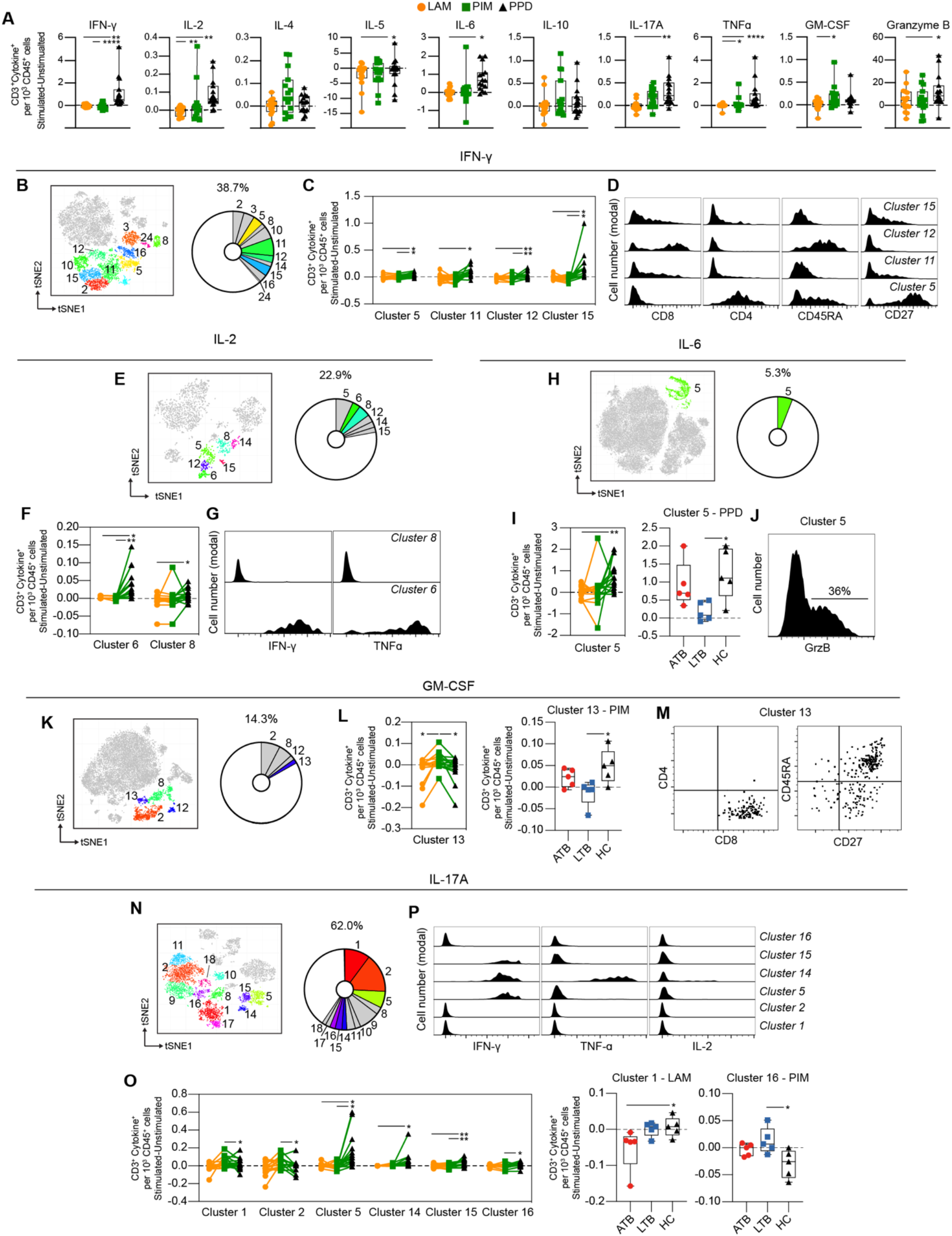
Cytokine production by stimulated T cells. (**A**) The number of cytokine-producing T cells per 1000 total CD45^+^ cells for each stimulation at 24 h with the background (unstimulated) cytokine-production removed. **(B)** Cluster analysis of IFN-*γ* secreting cells with clusters 2, 3, 5, 8, 10, 11, 12, 14, 15, 16, and 24 corresponding to T cells. **(C)** Clusters significantly affected by stimulation with **(D)** cluster histograms indicating CD4, CD8, CD45RA, and CD27. **(E)** Cluster analysis of IL-2 secreting cells with clusters 5, 6, 8, 12, 14, and 15 corresponding to T cells. **(F)** Clusters significantly affected by stimulation with **(G)** cluster histograms indicating IFN-*γ* and TNF-ɑ secretion. **(H)** Cluster analysis of IL-6 secreting cells with cluster 5 corresponding to T cells. **(I)** Cluster 5 is significantly affected by stimulation (left) and comparison of PPD stimulation on donors with active TB (ATB), latent TB (LTB) and healthy controls (HC) in cluster 5 (right). **(J)** Cluster histogram indicating GrzB secretion. **(K)** Cluster analysis of GM-CSF secreting cells with cluster 2, 8, 12, and 13 corresponding to T cells. **(L)** Cluster 13 is significantly affected by stimulation (left) and comparison of PIM stimulation on donors with ATB, LTB and HC in cluster 13 (right). **(M)** Cluster dot plots indicating CD4, CD8, CD45RA, and CD27 expression. **(N)** Cluster analysis of IL-17A secreting cells with cluster 1, 2, 5, 8, 9, 10, 11, 14, 15, 16, 17, and 18 corresponding to T cells. **(O)** Clusters significantly affected by stimulation (left) and comparison of PIM and LAM stimulation on donors with active ATB, LTB and HC in clusters 1 and 16, respectively (right) **(P)** Clusters histograms indicating IFN-*γ*, TNF-ɑ, and IL-2. Statistical differences between stimulations in (**A, I, and L**) were evaluated by Friedman’s test with Dunnet’s posttest, while comparisons within multiple clusters (**C, F, and O** left panels) were evaluated by a matched-pair two-way ANOVA with Geissner-Greenhouse correction followed by Tukey’s posttest (n=15/stimulation). Groups (ATB/LTB/HC) (**I, L, and O** right panels) were compared using Kruskal-Wallis with Dunn’s posttest (n=5/group) *p<0.05, **p<0.01, ***p<0.001, ****p<0.0001.

Approximately 23% of all IL-2 producing cells were identified as T cells (Figure 5E). These cells comprise six clusters, of which two (clusters 6 and 8) were significantly higher following PPD stimulation compared with LAM and/or PIM (Figure 5F). Cluster 6 corresponded to polyfunctional CD4^+^ T cells, co-producing IFN-γ and TNF-*α*, while cluster 8 was composed of cells producing only IL-2 (Figure 5G).

Although the regulatory effect of IL-6 on T cells is well known (29), the literature on IL-6 producing T cells is limited. We identified one cluster of IL-6^+^ T cells (cluster 5) corresponding to 5.3% of total IL-6^+^ leukocytes after 24 h stimulation (Figure 5H). This cluster was significantly elevated by PPD stimulation compared to LAM and was mainly attributed to ATB and HC, but not LTB individuals (Figure 5I). Cluster 5 was a mixed cluster consisting of cells expressing CD8^+^, CD4^+^, and double negative (DN) T cells (data not shown) with 36% of the cells also producing GrzB (Figure 5J).

Approximately 14% of the GM-CSF^+^ cells were T cells, represented by four different clusters (Fig 5K). Cluster 13 was significantly increased upon PIM stimulation compared to LAM and PPD (Figure 5L). The effect was more prominent in HC individuals compared to LTB. This cluster corresponded mostly to naïve (CD45RA^+^CD27^+^) CD8^+^ T cells (Figure 5M).

T cells represented 62% of the IL-17A^+^ cells (Figure 5N). Four out of 12 clusters (clusters 5, 14, 15, and 16) were increased by PPD compared with PIM and/or LAM stimulations, while clusters 1 and 2 were increased by PIM compared with PPD (Figure 5O left). In addition, the analysis of individual clusters showed that stimulation with LAM reduced cluster 1 in ATB, compared with HC individuals. Also, cluster 16 was higher in LTB compared with HC upon PIM stimulation (Figure 5O right). Three of these clusters corresponded to polyfunctional T cells (clusters 5, 14, and 15), with clusters 5 and 15 co-producing IFN-γ, and cluster 14 co-producing IFN-γ and TNF-*α* (Figure 5P).

In summary, T cell responses were mainly observed upon stimulation with PPD. The T cells producing IFN-γ, IL-2, IL-6, and IL-17A, some of those with a polyfunctional phenotype, were significantly increased with PPD compared with LAM and/or PIM. Interestingly, however, PIM stimulation led to an increase in GM-CSF-producing T cells, particularly in HC individuals, potentially indicating a different mechanism for GM-CSF induction also associated with disease status.

### NK cells are primarily stimulated by PPD

As for T cells, the NK cells (identified as CD3^−^CD7^+^) showed minor responses to PIM and LAM, and were mainly affected by PPD stimulation, which resulted in significantly higher numbers of NK cells producing IFN-γ, IL-2, IL-6, IL-17A, and GM-CSF, compared to PIM and LAM (Figure 6A). Most of IFN-γ producing cells at 24 h of stimulation are NK cells. They represented 51% of all IFN-γ-producing cells and could be further divided into 9 clusters (clusters 1, 6, 7, 13, 19, 21, 22, 23, and 25) (Fig 6B). Of these, four clusters were significantly increased following PPD stimulation, compared with LAM and PIM (Fig 6C). All of these clusters were CD57^−^ but expressed different levels of CD56 suggesting that they belonged to different NK subsets. Moreover, all clusters expressed GrzB while cluster 13 also expressed IL-17A (Fig 6D).

**Figure 6.**
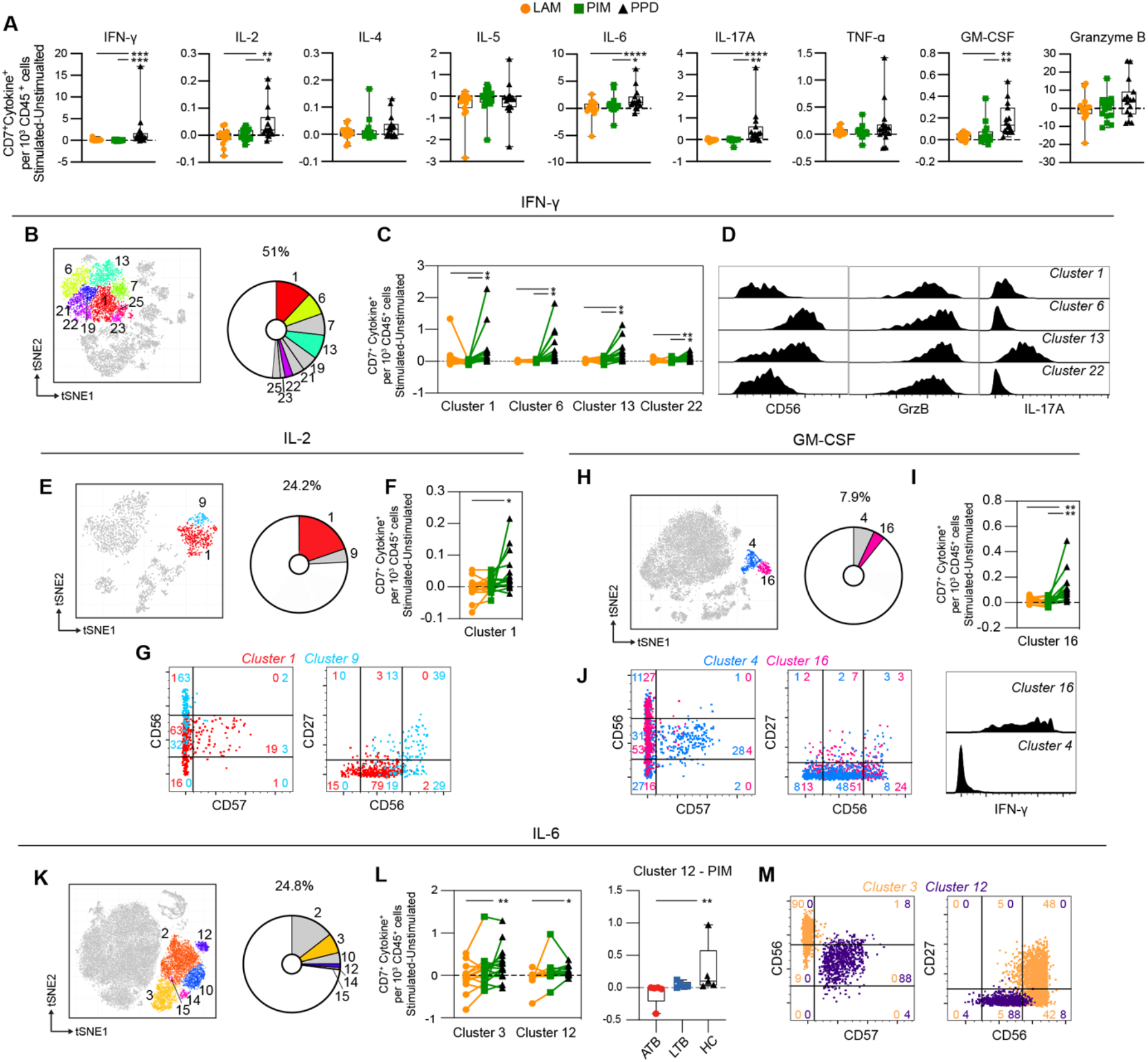
Cytokine production by stimulated NK cells. (**A**) The number of cytokine-producing NK cells per 1000 total CD45^+^ cells for each stimulation at 24 h with the background (unstimulated) cytokine-production removed. **(B)** Cluster analysis of IFN-*γ* secreting cells with clusters 1, 6, 7, 13, 19, 21, 22, 23, and 25 corresponding to NK cells. **(C)** Clusters significantly affected by stimulation with **(D)** cluster histograms indicating CD56 expression and GrzB and IL-17A secretion. **(E)** Cluster analysis of IL-2 secreting cells with cluster 1 and 9 corresponding to NK cells. **(F)** Clusters significantly affected by stimulation with **(G)** cluster dot plots showing CD56, CD57, and CD27 expression and histograms indicating IL-6 production. **(H)** Cluster analysis of GM-CSF secreting cells with cluster 4 and 16 corresponding to NK cells. **(I)** Cluster 16 significantly affected by stimulation. **(J)** Cluster’s dot plots showing CD56, CD57, and CD27 expression and histogram indicating IFN-*γ* secretion. **(K)** Cluster analysis of IL-6 secreting cells with cluster 1 and 9 corresponding to NK cells. **(L)** Clusters significantly affected by stimulation (left) and comparison of PIM stimulation on donors with active TB (ATB), latent TB (LTB) and healthy controls (HC) in cluster 12 (right). **(M)** Cluster’s dot plots showing CD56, CD57, and CD27 expression. Numbers in dot plots indicate the percentage within the cluster. Statistical differences between stimulations in (**A, F, and I**) were evaluated by Friedman’s test with Dunnet’s posttest, comparisons within multiple clusters (**C, L** left) were evaluated by a matched-pair two-way ANOVA with Geissner-Greenhouse correction followed by Tukey’s posttest (n=15/stimulation). Groups (ATB/LTB/HC) (**L**, right) were compared using Kruskal-Wallis with Dunn’s posttest (n=5/group) *p<0.05, **p<0.01, ***p<0.001, ****p<0.0001.

IL-2-producing NK cells constituted 24% of all IL-2^+^ cells and represented two clusters (cluster 1 and 9) (Fig 6E). Cluster 1 was significantly increased by PPD, compared with LAM stimulation (Fig 6F). Both clusters were CD57^−^ while cluster 1 expressed intermediate levels of CD56 and no CD27 while cluster 9 expressed high levels of CD56 and CD27 (Fig 6G). Both clusters co-produced IL-6 (Fig 6G). NK cells represent approximately 8% of all GM-CSF-producing cells at 24 h of stimulation (Fig 6H). The cytokine was produced by two clusters (4 and 16), one of which (cluster 16) was significantly higher in response to PPD compared with LAM and PIM stimulation (Fig 6I). Similar to PPD-mediated IL-2 secreting NK cells, GM-CSF was primarily produced by CD57^−^ NK cells where >50% expressed intermediate CD56 levels while almost no cells expressed CD27 (Fig 6J). Cluster 16 cells were also co-producing IFN-γ (Fig 6J). Approximately 25% of all IL-6-producing cells at 24h were identified as NK cells (Fig 6K), and two out of the six clusters (3 and 12) were significantly increased in numbers by PPD stimulation compared to LAM (Fig 6L). These two clusters belonged to different NK subsets with cluster 3 corresponding to CD56^high^CD57^−^ NK cells, of which 48% also expressed CD27, while cluster 12 was composed of CD56^int^CD57^+^CD27^−^ NK cells (Fig 6M). The IL-17A producing NK cells were composed of two clusters at 24h. However, they were not significantly different between the different stimulations (data not shown).

Thus, similar to T cells, NK cells were primarily induced to secrete cytokines through stimulation with PPD compared with the Mtb glycolipids LAM and PIM. The stimulation led to cytokine production by CD56^int^ and CD56^bright^ NK cells, independent on the expression of CD57. In summary, these results show that stimulation with PPD leads to rapid activation of different NK cell subsets with production of primarily pro-inflammatory cytokines.

### Atypical B cells are a major source of polyfunctional cytokine responses following PIM stimulation

Compared with T cells and myeloid cells, B cells (defined as CD3^−^HLA-DR^+^CD20^+^) were minor producers of the measured cytokines (Figure 3). There was however a primarily PIM-derived effect leading to significantly increased numbers of IL-4, IL-10, and GM-CSF producing B cells in comparison to LAM and/or PPD stimulation (Figure 7A). There were two B cell clusters producing IL-4 (cluster 6 and 12) (Figure 7B), but only cluster 6 was significantly increased by PIM stimulation, with PPD leading to the lowest numbers of cells in this cluster (Figure 7C). Cluster 6 was enriched for switched memory (CD27^+^IgD^−^) and double negative (DN; CD27^−^IgD^−^) B cells, while cluster 12 was enriched for naïve B cells (CD27^−^IgD^+^) (Figure 7D). Cluster 6 was enriched for CD11c^+^ B cells, which are associated with recent B cell activation and formation of atypical B cells during infection or inflammatory conditions (30).

**Figure 7.**
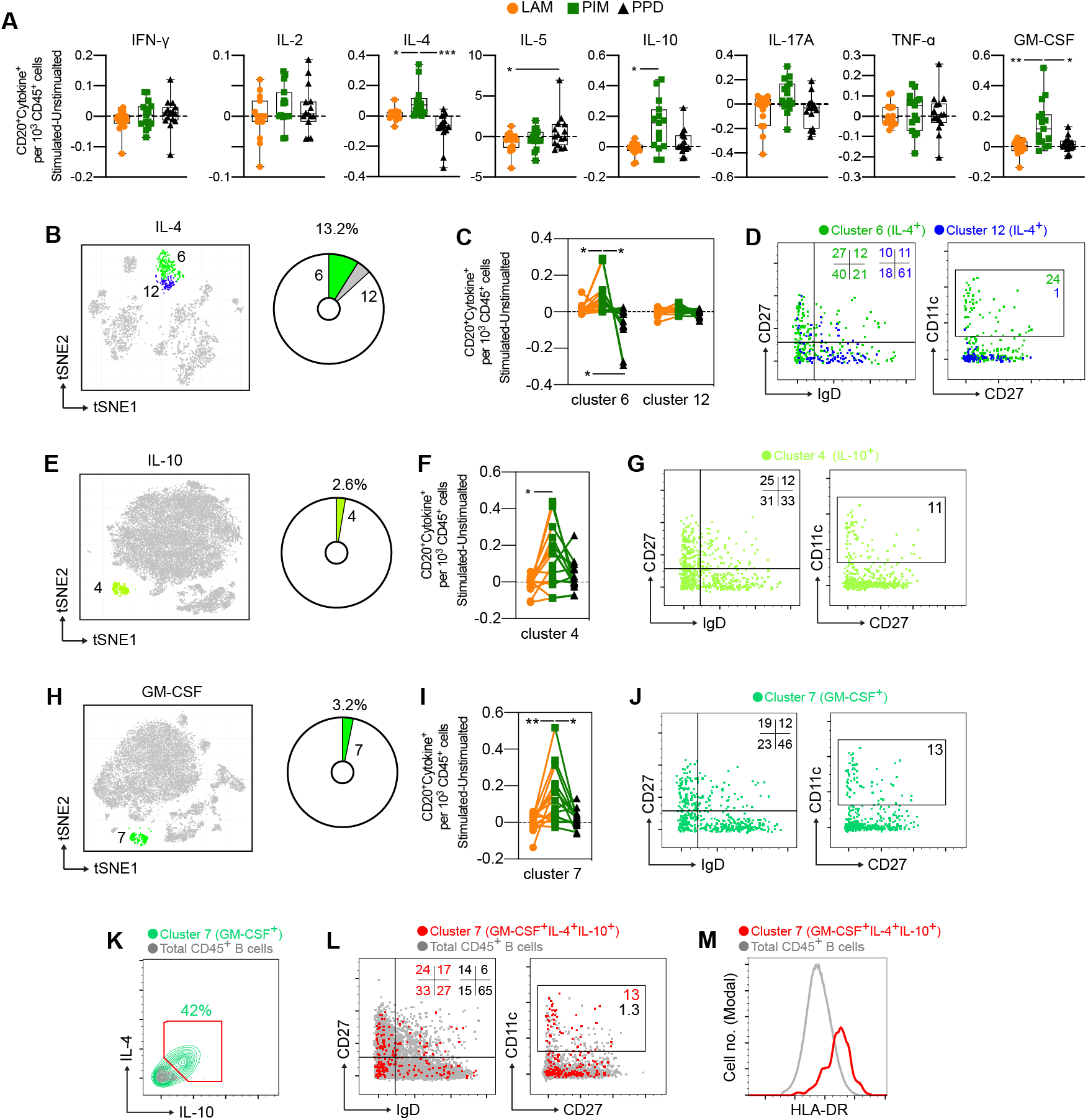
Cytokine production by stimulated B cells. (**A**) The number of cytokine positive B cells per 1000 total CD45^+^ cells for each stimulation at 24 h with the background (unstimulated) cytokine-production removed. (**B**) Cluster analysis of IL-4 secreting cells with cluster 6 and 12 corresponding to B cells. The pie-chart indicates cluster-specific and percent of total contribution to all IL-4 secreting cells. (**C**) Evaluation of the effect of LAM (orange circles), PIM (green boxes), or PPD (black triangles) stimulation on IL-4 secreting B cell clusters. (**D**) Overlay of concatenated IL-4 secreting B cells for cluster 6 (green) and cluster 12 (blue) assessing IgD and CD27 or CD11c and CD27 surface expression. (**E-G**) A similar analysis for IL-10 secreting cells and (**H-J**) GM-CSF secreting cells. (**K**) IL-4 and IL-10 co-expression among GM-CSF^+^ B cells (green) and total B cells (grey). (**L**) Overlay scatter plot of GM-CSF^+^IL-4^+^IL-10^+^ triple-secreting B cells (red) and total B cells (grey) assessing IgD and CD27 or CD11c and CD27 surface expression, with frequency of cells included in the gates indicated. (**M**) Overlay histogram indicating HLA-DR expression for GM-CSF^+^IL-4^+^IL-10^+^ triple-secreting B cells (red) and total B cells (grey). Numbers in dot plots indicate the percentage within the cluster. Statistical differences between stimulations in individual groups (**A, F, I**) were evaluated by Friedman’s test with Dunnet’s posttest, while comparisons within multiple clusters (**C**) were evaluated by a matched-pair two-way ANOVA with Geissner-Greenhouse correction followed by Tukey’s posttest (n=15/stimulation). *p<0.05, **p<0.01, ***p<0.001. n=15 for each group. Scatter and overlay plots show data concatenated from all samples and donors (n=60).

B cells producing IL-10 and GM-CSF were also significantly expanded by PIM stimulation (Figure 7E-J). As B cells responding to PIM stimulation presented a highly homogenous phenotype, we further evaluated the cells for co-expression of the three cytokines (Figure 7K). We found that 42% of GM-CSF-producing B cells also produced IL-4 and IL-10. Compared with total B cell populations, the phenotype of the polyfunctional cells was highly enriched for double negative (DN - IgD^−^CD27^−^) B cells but also for unswitched and switched memory B cells (CD27^+^) compared with total B cell populations (Figure 7L). The polyfunctional B cells were also approximately 10-fold enriched for CD11c^+^ B cells compared with total B cells, suggesting that atypical B cells can respond to PIM stimulation (Figure 7L). We also quantified the levels of HLA-DR on the cell surface of the polyfunctional B cells and compared with the levels on total B cells and found an increased expression of HLA-DR on cells from cluster 7 (Figure 7M), consistent with previous reports on atypical B cells in mice and humans (31, 32).

### Rapid polyfunctional response of myeloid cells to PIM stimulation

The production of cytokines by CD33^+^ myeloid cells was compared for each stimulation (Figure 8A). PIM stimulation led to a robust increase of cells producing IL-2, IL-4, IL-6, IL-10, TNF-*α*, compared to PPD, and of IL-6, IL-17A, TNF-*α* and GM-CSF in comparison to LAM (Figure 8A). Interestingly, IL-10 producing cells were induced by both PIM and PPD (Figure 8A), contrasting with the other cytokines that were primarily induced by PIM.

**Figure 8.**
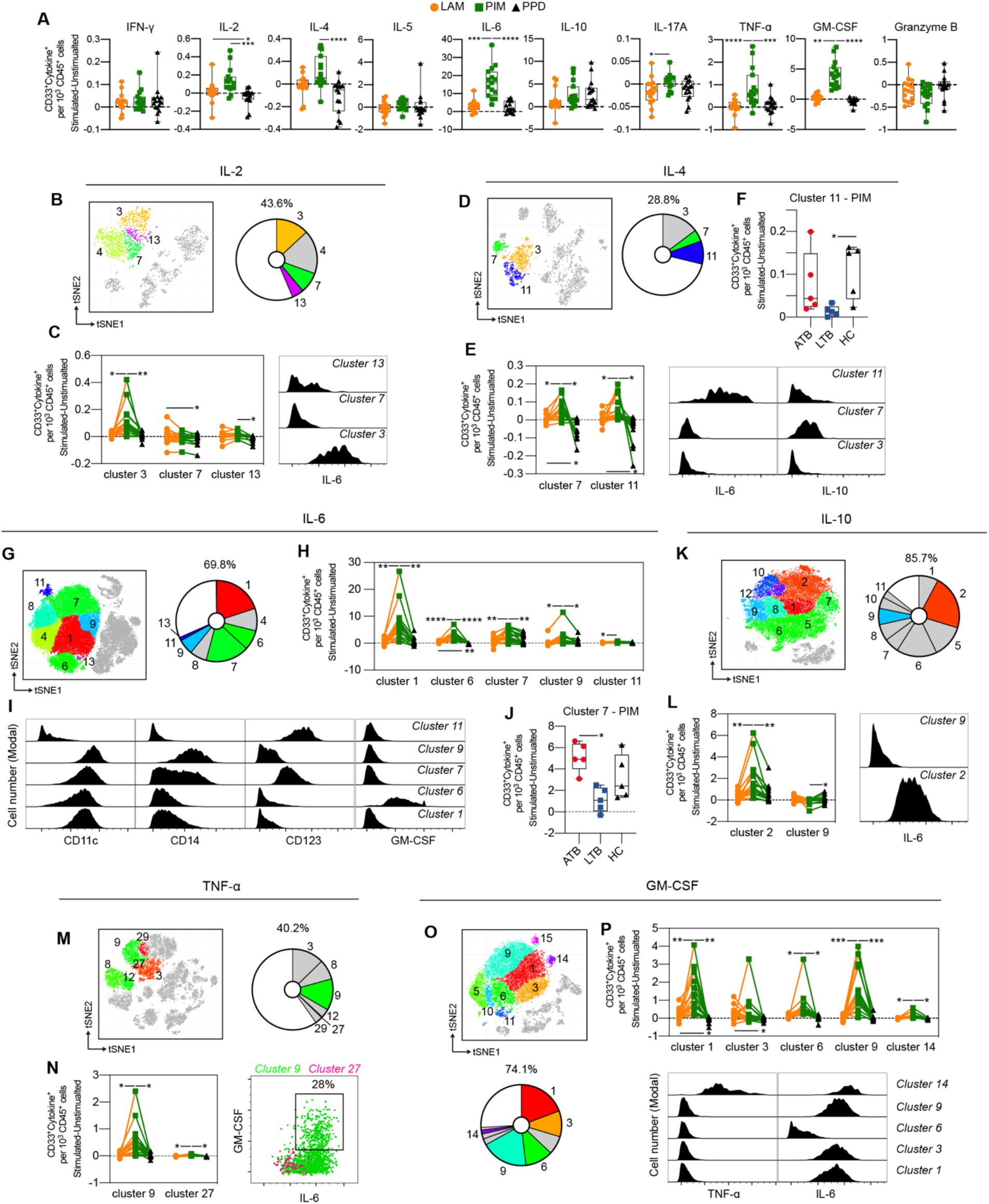
Cytokine production by stimulated CD33^+^ myeloid cells. (**A**) The number of cytokine positive CD33^+^ myeloid cells per 1000 total CD45^+^ cells for each stimulation at 24 h with the background (unstimulated) cytokine-production removed. (**B**) Cluster analysis of IL-2 secreting cells with cluster 3, 4, 7 and 13 corresponding to myeloid cells. (**C**) Clusters significantly affected by stimulation (left) with cluster histograms indicating co-secretion of IL-6. (**D**) Cluster analysis of IL-4 secreting with cluster 3, 7, and 11 corresponding to myeloid cells. (**E**) Clusters significantly affected by stimulation (left) and co-expression with IL-6 and IL-10 (right). (**F**) Comparison of PIM stimulation on donors with active TB (ATB), latent TB (LTB) and healthy controls (HC) in cluster 11. (**G**) Myeloid clusters secreting IL-6 (**H**) significantly affected by stimulation. (**I**) Cell surface phenotype of indicated cluster. (**J**) Differential effect of PIM stimulation on cluster 7 cells in ATB, LTB, and HC. (**K**) Myeloid clusters secreting IL-10 with (**L**) clusters significantly affected by stimulation (left panel) and histograms indicating IL-10 co-expression with IL-6. (**M**) Myeloid clusters secreting TNF⍰-ɑ (**N**) significantly affected by stimulation (left) with co-expression of GM-CSF and IL-6 (right). (**O**) Myeloid clusters secreting GM-CSF. (**P**) Clusters differently affected by stimulation (left) with co-expression of TNFa and IL-6 (right). Statistical differences between stimulations in (**A**) were evaluated by Friedman’s test with Dunnet’s posttest, while comparisons within multiple clusters (**C, E, H, L, N, P**) were evaluated by a matched-pair two-way ANOVA with Geissner-Greenhouse correction followed by Tukey’s posttest (n=15/stimulation). Groups (ATB/LTB/HC) (**F, J**) were compared using Kruskal-Wallis with Dunn’s posttest (n=5/group) *p<0.05, **p<0.01, ***p<0.001, ****p<0.0001.

To understand if the effect of stimulation was associated with specific myeloid subsets, we further investigated the impact of stimulation on individual cell clusters. Since monocytes loose CD16 expression during culture (33), it is difficult to distinguish non-classical monocytes from myeloid dendritic cells (mDCs), and intermediate and classical monocytes. In separate FACS experiments, we found, however, that CD123 (corresponding to the IL-3R), is commonly upregulated on non-classical monocytes and somewhat on intermediate monocytes (data not shown), enabling distinction of the different cell subsets. The IL-2 producing myeloid cells constituted 43.6% of all IL2-producing cells and were composed of four different clusters (cluster 3, 4, 7, and 13), of which three were differently affected by the stimuli (Fig 8B). For cluster 3 and 13, PIM stimulation led to significantly more IL-2^+^ cells compared with PPD and/or LAM, while cluster 7 was higher in LAM compared to PPD Fig 8C). Cluster 7 expressed CD14, while clusters 3 and 13 were mostly negative for CD14 (Supplemental Figure 5). Cluster 3 was associated with the co-production of IL-6 (Figure 8C).

Approximately 29% of all IL-4 producing cells after 24h of stimulation expressed CD33. (Figure 8D). These cells were further distributed into three clusters (3, 7, and 11), of which cluster 7 and 11 were significantly higher in number following PIM stimulation compared with LAM and PPD stimulation. LAM stimulation also led to more IL-4 producing cells compared to PPD (Figure 8E). Both cluster 7 and 11 produced several other cytokines in addition to IL-4, with cluster 7 also producing IL-10, and cluster 11 producing IL-6, and IL-10 (Figure 8E). Interestingly, this effect of multiple cytokine production, was significantly reduced in individuals with LTB compared with ATB and HC (Figure 8F).

IL-6 was the most frequent cytokine produced following PIM stimulation (Fig 8A). Approximately 70% of all IL-6 secreting cells at 24h were myeloid cells (Fig 5G) with 5 out of 7 clusters showing a significant increase following PIM stimulation compared with PPD and/or LAM (Fig 8H). Several subsets of myeloid cells responded with IL-6 production, including CD14^−^ CD123^−^ DCs (cluster 1 and 6), intermediate/non-classical monocytes (CD14^int/–^CD123^+^, cluster 7), and classical monocytes (CD14^+^CD123^−^, cluster 9). Among these, the cluster 6 DCs, in addition to IL-6, also produced GM-CSF (Figure 8I). Similar to the IL-4^+^IL-6^+^ co-producing cluster 11 (Figure 8F), the intermediate/non-classical monocyte cluster 7 contracted in individuals with LTB, compared with those with ATB and HC (Figure 8J).

Myeloid cells were the main cell subset identified within IL-10, TNF*α* and GM-CSF-producing cells, especially following stimulation with PIM (Figure 8K, M, O). One IL-10 clusters, two TNF-*α* clusters and five GM-CSF clusters were significantly increased in comparison to LAM and PPD (Figure 8L, N, P). Of these, parts of TNF-*α* cluster 9 and GM-CSF cluster 14 likely corresponded to the same polyfunctional cells as both clusters secreted TNF-*α*, GM-CSF, and IL-6 (Figure 8L, N), while one of the two IL-10 clusters that were affected by PIM co-produced IL-6 only, the other only produced IL-10. GM-CSF cluster 1, 3, and 9 also co-produced IL-6, but not TNF*α*, while cluster 6 only produced GM-CSF.

In summary, several myeloid cell subsets rapidly responded to stimulation by producing cytokines. The response was primarily induced by PIM and included phenotypes of cells producing both single and multiple cytokines. Among the most polyfunctional responses were cells producing IL-4, IL-6, and IL-10, or TNF-*α*, GM-CSF and IL-6.

### Stimulation of myeloid cells with PIM is partially TLR2 dependent

PIM and LAM stimulation induced a robust immune response in myeloid cells (Figure 8A). To investigate the mechanism responsible for this effect, and in particular the dependence on interaction with TLR2, PBMCs from HC were treated with anti-TLR2 blocking antibody before stimulation with PIM, LAM and PHA. Blocking TLR2 led to a reduction in the percentage of CD33^+^IL-6^+^ myeloid cells upon PIM stimulation, but not with LAM or PHA (Fig 9). We did not observe any significant effect of blocking TLR2 on IL-6 production from T cells, NK cells, or B cells (data not shown), although the frequency of IL-6^+^ cells was very low on those cell subsets. Our results show that PIM engagement of TLR2 contributes to IL-6 production in myeloid cells, although as the blocking effect was not complete, other mechanisms of PIM stimulation likely remains.

**Figure 9.**
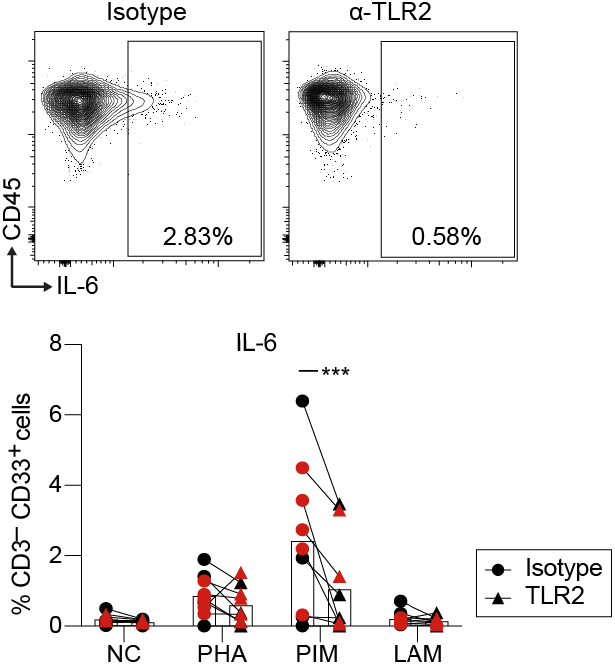
IL-6 production by myeloid cells upon PIM stimulation is regulated via TLR2. Dot plots of IL-6 gating within myeloid cells (top) and percentage of cells producing IL-6 within myeloid cells (bottom). PBMCs were pre-treated with anti-TLR2 or isotype control and then stimulated with PIM, LAM, and PHA during 24h. IL-6 production was evaluated by flow cytometry. Statistical differences between anti-TLR2 and isotype were evaluated by a paired t test (n=9) ***p<0.001. NC = negative control. The different colors correspond to two independent experiments.

## Discussion

In the present study, we used mass cytometry and secretome analysis to assess the effect of stimulation with two mycobacterial glycolipids on PBMCs from individuals with ATB or LTB. We show that LAM and PIM induce responses in PBMCs from Mtb-infected individuals that are quite distinct from those obtained from HC. In addition, we show that the responses to these glycolipids are clearly different from those elicited by PPD. The responses involve both expansion and contraction of particular cell subsets, production and secretion of distinct patterns of cytokines and chemokines.

When staining for intracellular cytokine production, we found that PIM mainly induced antigen-presenting cells to produce a defined set of pro-inflammatory cytokines consisting of IL-2, IL-6, IL-17A, TNF-*α* and GM-CSF, the anti-inflammatory IL-10 as well as IL-4, but not IFN-γ. LAM triggered responses that tended to be similar to the ones generated by PIM, but weaker in most instances. Classical and intermediate monocytes are known to secrete high levels of pro-inflammatory cytokines in response to microbial products (34). In addition, compared to non-classical monocytes, they were previously shown to exhibit a greater polyfunctional pro-inflammatory response (IL1-α, IL1-β, IL-6, IL-8, IL-10, and TNF-*α*) to lipomannan from *Mycobacterium smegmatis* (TLR-2 agonist) and LPS (TLR-4 agonist) (34). Here we show that PIM induced multifunctional monocytes producing cytokines in a combination of either pro-inflammatory IL-2, IL-6, GM-CSF and TNF-α, or IL-4 and the anti-inflammatory IL-10. In particular GM-CSF, which is increasingly recognized for its potential role in innate resistance to TB (35), was in our study mainly produced by myeloid cells upon PIM stimulation.

This response contrasted with the quite well-known immune response triggered by PPD, which was dominated primarily by T and NK cells. They produced predominantly the pro-inflammatory cytokines IFN-γ, IL-2, IL-6, IL-17A, TNF-*α*, and GrzB, but also IL-10, although no IL-4. While T cells simultaneously producing combinations of cytokines have been extensively investigated in the context of the immune response in TB (36-38), we extended these findings to several other cell types. Our results reveal that multiple subsets of myeloid cells, NK, B and T cells respond to glycolipids and/or to PPD, with the production of different combinations of cytokines such as classical functional T cells producing IFN-γ, IL-2 and TNF-*α*, but also other combinations, such as IL6 and GrzB or IL-17A and TNF-*α* with or without IFN-γ.

B cells producing IL-10 and GM-CSF are known to be present at relatively low frequencies in human peripheral blood (39, 40). This is in agreement with our finding that B cells were minor producers of IL-10 and GM-CSF, even after stimulation. We did however identify subsets of polyfunctional B cells that produced a combination of GM-CSF, IL-4 and IL-10. These cells were enriched among DN (CD27^−^IgD^−^) B cells and unswitched and switched memory B cells (CD27^+^). The polyfunctional B cells were also approximately 10-fold enriched among B cells expressing CD11c, which was recently associated with B cell activation and formation of atypical B cells (30), also known to expand during ATB (41). The function of these cells remains controversial with evidence of pro-inflammatory properties but also an association with B cell dysfunction due to reduced responsiveness following antigen binding (42). In mice a similar CD11c-expressing B cell subset was associated with improved APC function (31), which is supported by a recent report showing that CD21^lo^ B cells display improved T cell activation in vitro (32). Our findings in the present study would further support such a function as the polyfunctional B cells display elevated levels of HLA-DR combined with polarizing and immunomodulatory cytokines, important for T cell regulation. Human GM-CSF–expressing B cells are notable for being among the highest producers of both TNF-α and IL-6, and most *in vitro*-induced human IL-10^+^ B cells are also reported to secrete TNF-α and/or IL-6 (43). However, human B cell subsets have been reported to show a near-mutually exclusive expression of GM-CSF and IL-10 (39). By contrast, in our study, B cells stimulated by PIM did not co-produce GM-CSF with TNF-α or IL-6, but rather with IL-4 and IL-10, indicating a different pathway of stimulation.

We identified several polyfunctional T cell subsets producing combinations of IFN-γ/IL-2/IL-6/TNF-α/IL-17A that were expanded by PPD stimulation. Polyfunctional Mtb-specific T cells producing IFN-γ in combination with IL-2 and TNF-*α* were initially considered a surrogate of protection, in particular in vaccination studies against TB (36, 37, 44). However, this is a controversial issue (38) and other studies suggest that these cells are simply associated with active disease and not with protection (45, 46). Stimulation with PPD or Mtb protein/peptides has been reported to induce different T cell cytokine patterns in patients with ATB and LTB (45, 47). Compared to patients with ATB, individuals with LTB have been previously shown to have significantly lower proportions of Mtb-specific T cells secreting only IFN-γ and significantly higher proportions of cells secreting IFN-γ/IL-2 or just IL-2 (47). Mtb-specific CD4^+^ T cells expressing IFN-γ/IL-2/TNF-α were previously detectable in 85-90% of patients with ATB, but only 10-15% of subjects with LTB (45). On the contrary, LTB subjects had significantly higher (12-to 15-fold) proportions of IFN-γ/IL-2 double and IFN-γ single expressors as compared with the other CD4^+^ T cell subsets (45). We did not see significant differences in simultaneous IFN-γ/IL-2/TNF-α production upon PPD stimulation between groups although there was a tendency of less production of total cytokines in LTB. In contrast to PPD stimulation, PIM and LAM did not trigger detectable polyfunctional T cells

One interesting observation in our study is that upon stimulation with PIM, LAM or PPD, cells and supernatants from individuals with ATB or LTB produce less cytokines than the cells from HC. This is obvious for myeloid cells and to a lesser extent for B and T cells. The hyporesponsive state in monocytes in response to PIM and LAM is compatible with trained immunity leading to a tolerogenic cellular response. Trained immunity is defined as a long-term adaptation of innate immune cells leading either to an enhanced responsiveness or a tolerance state to a subsequent challenge (48, 49). In both cases, an initial stimulus induces epigenetic and metabolic changes in innate immune cells that result in a stronger or weaker response — training or tolerance, respectively — to a subsequent challenge that occurs days to months later. Different stimuli (for example, β-glucan, LPS, BCG/PPD) can induce different trained immunity programs (49). This type of adaptive features of innate immunity has been demonstrated in multiple studies of memory phenotypes in monocytes and Mφs (50-52). Chronic or repeated stimulation through TLRs can render immune cells unresponsive to subsequent challenges with the same or different TLR ligands (53, 54) as reviewed in (55). Stimulation of human monocytes with TLRs in high doses, with the exception of TLR9, consistently induce long-term tolerance to later stimulation with bacterial ligands (56). Our results support the hypothesis that repeated stimulation with LAM, and specifically with PIM, in ATB and LTB individuals lead to a reduced response to these molecules compared with HC.

The response of myeloid cells to PPD was weaker in LTB compared to HC, indicating hyporesponsiveness also to PPD. This is in line with earlier observations of depression of PPD-induced proliferative responses by monocytes from TB patients (57, 58), where direct stimulation of monocytes primed during TB infection appear to be responsible for *in vitro* suppression of PPD responses (57). Interestingly we also found that T cells were hyporesponsive to PIM. The overall cytokine production was reduced in individuals with ATB upon PIM stimulation. These results are in agreement with the systematic review by Li et al that found higher levels of IL-17 and IFN-*γ* in LTB when compared to ATB (59). An additional interesting observation in our study was the clear ability of PIM to expand GMCSF^+^ CD8^+^ T cells in HC but not in LTB patients. This hyporesponsiveness to PIM might be caused by T cell exhaustion or tolerance in Mtb infected individuals. Exhaustion of T cells represents a state of functional hyporesponsiveness due to persistent antigen exposure and inflammation reported for TB and other chronic infections (60-62). In mice this effect has also been shown upon repeated exposure to mycobacterial antigens (63), including LAM (64). The hyporesponsiveness of T cells to PIM in our study could be the result of direct or indirect inhibition. Antigen-specific CD4^+^ T-cell activation can be directly inhibited by LAM (65-68) and PIM (66). By interfering with very early events in TCR signaling, LAM and PIM may drive cells to a state of anergy (66, 68), which could provide another explanation of the poor response of cells from ATB and LTB individuals to Mtb glycolipids. Alternatively, the hyporesponsiveness could be indirect, through upstream effects of hyporesponsive myeloid cells, since PIM and LAM also induce proliferation of specific T cells upon presentation by CD1 molecules on myeloid cells.

IL-6 is known to be strongly induced in monocytes and DCs upon TLR2 ligation (69). We observed that PIM stimulation induced IL-6 production mainly in myeloid cells (DCs and classical/nonclassical monocytes). Moreover, treatment with an anti-TLR2 antibody led to partial inhibition of PIM-induced IL-6 production in myeloid cells, suggesting that PIM induces IL-6 production through TLR2. This is in line with other studies where it was observed that PIMs and ManLAM from Mtb induce pro-inflammatory cytokine production in human and mouse Mφs *via* recognition by TLR2 (70-72). However, IL-6 production was not completely abolished suggesting that other receptors could also be involved.

LAM and PIM had very similar effects on monocytes/DCs and T cells, although LAM induced a weaker response than PIM. Presuming that LAM and PIM act through the same TLR2 pathway the different responses are potentially associated with structural differences, where a common active site may be partly masked in LAM compared to PIM. Nigou et al showed that LAM induces a weaker signal through TLR2 compared to PIM_6_, suggesting that the bulky arabinan domain may mask the mannan chain in such a way that they behave like molecules with a mannan restricted to a single mannosyl unit (72). This is also in line with observations by Shukla et al. that PIM_6_ induces TLR2-mediated extracellular-signal-regulated kinase (ERK) activation and TNF-*α* secretion in Mfs, while LAM was not an effective functional activator of TLR2 signaling (73). The weaker effect of LAM compared to PIM may also in part depend on the fact that the LAM that was used in the present study has a much higher molecular weight compared to PIM resulting in a lower molar concentration.

In contrast to the glycolipids, PPD displayed a markedly different response, mainly by inducing IFN-*γ*. PPD contains a complex mixture of proteins, including the antigens ESAT-6 and CFP10 that are the antigens used in the Mtb specific IFN-*γ* release assays (IGRA-tests). We did not identify which antigens in PPD that were responsible for the immune responses presented in this study. However, since PPD is still widely used in clinical testing, the unprecedented level of details presented here may be useful to better understand how individual immune cell subsets react with Mtb proteins.

In conclusion, Mtb glycolipids induced a T_H_2/T_H_17-biased immune response, partially by strongly affecting the myeloid cells, while PPD primarily induced a T_H_1 response mostly mediated by T and NK cells. It is generally thought that a balance between T_H_1 and T_H_17 responses is needed to control bacterial growth and limit immunopathology during TB. A shift towards exaggerated IL-17A production has been associated with prolonged neutrophil recruitment and tissue damage (74, 75). Our results suggest that the response to PPD (promoting IFN-γ producing cells) is counterbalanced by the response to PIM that rather promotes IL-17A producing cells.

The hyporesponsive state of monocytes observed in ATB and LTB in response to PIM was more prominent in LTB. The immune profile in LTB is thought to represent a more protective pattern than in ATB (76, 77). It is possible that during LTB a continuous level of stimulation maintains a pool of protective memory cells (18), while at the same time inducing tolerance in monocytes, which could indicate protection of the host from excessive production of pro-inflammatory cytokines and control of lung tissue damage (78). By defining the mechanistic background of the response to Mtb glycolipids such as PIM, that combine the induction of innate and adaptive immune memory, new vaccine strategies and correlates of protection may be developed.

## Acknowledgements

We thank all study participants that were recruited into the study. We also thank Julius Lautenbach for critically reading the manuscript. The work presented was performed at Karolinska Institutet and Life and Health Sciences Research Institute (ICVS), University of Minho. Financial support was provided by grants from the Swedish Research Council (grant 2016-05683 and 2020-03602) and the Swedish Heart-Lung Foundation (grants 20160336, 20180386, and 20200194) to GK. Grants from Clas Groschinsky’s memorial foundation (M2049), Åke Wiberg’s foundation (M19-0559), the Swedish Medical Association (SLS-934363), and Magnus Bergvall’s foundation (2019-03436) to CS. Grants from the Foundation for Science and Technology (FCT) - project UIDB/50026/2020 and UIDP/50026/2020 to MCN. CSS is supported by an FCT PhD grant, in the context of the Doctoral Program in Applied Health Sciences (PD/BDE/142976/2018). JC-G is supported by an FCT PhD grant, in the context of the Doctoral Program in Aging and Chronic Diseases (PD/BD/137433/2018). CN is a junior researcher under the scope of the FCT Transitional Rule DL57/2016. We would like to thank the team from the Translational Plasma Profile Facility at SciLifeLab for support and the generation of data for this project.

## Author contributions

GK, MC-N, and CS designed the study. CSS, CS, CN, JC-G, and TL performed experiments and/or analysis. EF, GF, and JB included patients and provided clinical data. CSS, CS, and EF generated the figures and tables. BC and PB provided key resources. CSS, CS, EF, MC-N, and GK wrote the first draft and all authors contributed to manuscript revision.

