## Supplemental material for "Stimulation with mycobacterial glycolipids and PPD reveals different innate immune response profiles in active and latent TB"

**Supplemental Table 1. Study individuals**

| | ATB (n=5) | LTBI (n=5) | HC (n=5) |
|---|---|---|---|
| **Male sex** | 5 | 2 | 4 |
| **Age (median, range)** | 39 (24-50) | 39 (25-72) | 25 (22-37) |
| **Comorbidity** | 2* | 0 | 0 |
| **Born in TB high endemic country** | 4 | 3 | 0 |
| **Years since immigration to Sweden** | 8 (0.5-20) | 11 (3-16) | NA |
| **BCG vaccination** | unknown | 3 | 1 |
| **Symptoms** | | | |
| **- cough** | 4 | NA | NA |
| **- fever/night sweats/weight loss** | 3 | | |
| **IGRA** | | | |
| **- positive** | NA | 4 | 0 |
| **- conversion (after 2 months)** | | 1 | 0 |
| **- negative** | | 0 | 5 |
| **Chest X-Ray/Computed Tomography** | | | |
| **- infiltrates** | 5 | 2** | NA |
| **- cavitaries** | 2 | 0 | |
| **- pleural effusion** | 3 | 0 | |
| **Mycobacteriology** | | | |
| **- microscopy positive** | 1 | 0 | NA |
| **- PCR positive** | 1 | 0 | |
| **- culture positive** | 5 | 0 | |

*Diabetes mellitus type 2, chronic kidney disease stage 2; **Normal on repeat radiology

NA: not applicable; ATB: Active TB; LTB: latent TB; HC: Healthy controls.

**Supplemental Table 2. Broad staining panel for mass cytometry.**

| Marker | Clone | Tag | Company | Dilution |
|---|---|---|---|---|
| CD45 | HI30 | 89Y | Fluidigm | 1:200 |
| CD57 | HCD57 | 115In | BioLegend | 1:100 |
| CD19 | HIB19 | 142Nd | BioLegend | 1:100 |
| CD5 | UCHT2 | 143Nd | BioLegend | 1:200 |
| CD16 | 3G8 | 144Nd | BioLegend | 1:100 |
| CD4 | RPA-T4 | 145Nd | BioLegend | 1:100 |
| CD11c | Bu15 | 147Sm | Fluidigm | 1:100 |
| CD123 | 6H6 | 151Eu | BioLegend | 1:100 |
| CD3e | UCHT1 | 154Sm | Fluidigm | 1:200 |
| CD14 | M5E2 | 160Gd | BioLegend | 1:100 |
| CD161 | HP-3G10 | 161Dy | BioLegend | 1:100 |
| CD127 | A019D5 | 165Ho | Fluidigm | 1:100 |
| CD38 | HIT2 | 168Er | BioLegend | 1:100 |
| CD45RA | HI100 | 169Tm | Fluidigm | 1:100 |
| CD20 | 2H7 | 170Er | BioLegend | 1:100 |
| IgD | IA6-2 | 172Yb | BioLegend | 1:100 |
| Cell-ID™ Intercalator-Ir | DNAIr-191 | - | Fluidigm | 1:1000 |
| Cell-ID™ Intercalator-Ir | DNAIr-193 | - | Fluidigm | 1:1000 |
| CD31 | WM59 | 148Nd | BioLegend | 1:200 |
| HLA-DR | L243 | 163Dy | BioLegend | 1:200 |
| CD44 | BJ18 | 174Yb | BioLegend | 1:100 |
| CD8a | SK1 | 146Nd | BioLegend | 1:300 |
| CD11b (MAC1) | Mac-1 | 209Bi | Fluidigm | 1:200 |
| CD56 | NCAM16.2 | 173Yb | BD | 1:200 |
| CD7 | CD7-687 | 155Gd | BioLegend | 1:200 |
| CD27 | L128 | 167Er | Fluidigm | 1:200 |
| Siglec-8 | 7C9 | 164Dy | Fluidigm | 1:100 |
| CD33 | WM53 | 157Gd | BioLegend | 1:100 |
| TCRγδ | 5A6.E9 | 152Sm | Fisher | 1:100 |

**Supplemental Table 3. Intracellular staining panel for mass cytometry.**

| Marker | Clone | Tag | Company | Dilution |
|---|---|---|---|---|
| IL-2 | MQ1-17H12 | 158Gd | Fluidigm | 1:200 |
| IL-4 | MP4-25D2 | 149Sm | BioLegend | 1:100 |
| IL-5 | TRFK5 | 153Eu | BioLegend | 1:100 |
| IL-6 | MQ2-13AS | 156Gd | Fluidigm | 1:100 |
| IL-10 | JES3-9D7 | 159Tb | BioLegend | 1:100 |
| IL-17A | N49-653 | 166Er | BioLegend | 1:100 |
| IFN-γ | B27 | 150Nd | BioLegend | 1:125 |
| TNF | MAb11 | 175Lu | BioLegend | 1:75 |
| Granzyme B | GB11 | 171Yb | Fluidigm | 1:100 |
| GM-CSF | BVD2-21C11 | 176Yb | BioLegend | 1:75 |

**Supplemental Table 4. Olink inflammation panel.**

| Protein | UniProt ID | Protein | UniProt ID |
|---|---|---|---|
| Adenosine Deaminase (ADA) | P00813 | Fractalkine (CX3CL1) | P78423 |
| Artemin (ARTN) | Q5T4W7 | Glial cell line-derived neurotrophic factor (GDNF) | P39905 |
| Axin-1 (AXIN1) | O15169 | Hepatocyte growth factor (HGF) | P14210 |
| Beta-nerve growth factor (Beta-NGF) | P01138 | Interferon gamma (IFN-gamma) | P01579 |
| Caspase-8 (CASP-8) | Q14790 | Interleukin-1 alpha (IL-1 alpha) | P01583 |
| C-C motif chemokine 3 (CCL3) | P10147 | Interleukin-2 (IL-2) | P60568 |
| C-C motif chemokine 4 (CCL4) | P13236 | Interleukin-2 receptor subunit beta (IL-2RB) | P14784 |
| C-C motif chemokine 19 (CCL19) | Q99731 | Interleukin-4 (IL-4) | P05112 |
| C-C motif chemokine 20 (CCL20) | P78556 | Interleukin-5 (IL5) | P05113 |
| C-C motif chemokine 23 (CCL23) | P55773 | Interleukin-6 (IL6) | P05231 |
| C-C motif chemokine 25 (CCL25) | O15444 | Interleukin-7 (IL-7) | P13232 |
| C-C motif chemokine 28 (CCL28) | Q9NRJ3 | Interleukin-8 (IL-8) | P10145 |
| CD40L receptor (CD40) | P25942 | Interleukin-10 (IL10) | P22301 |
| CUB domain-containing protein 1 (CDCP1) | Q9H5V8 | Interleukin-10 receptor subunit alpha (IL-10RA) | Q13651 |
| C-X-C motif chemokine 1 (CXCL1) | P09341 | Interleukin-10 receptor subunit beta (IL-10RB) | Q08334 |
| C-X-C motif chemokine 5 (CXCL5) | P42830 | Interleukin-12 subunit beta (IL-12B) | P29460 |
| C-X-C motif chemokine 6 (CXCL6) | P80162 | Interleukin-13 (IL-13) | P35225 |
| C-X-C motif chemokine 9 (CXCL9) | Q07325 | Interleukin-15 receptor subunit alpha (IL-15RA) | Q13261 |
| C-X-C motif chemokine 10 (CXCL10) | P02778 | Interleukin-17A (IL-17A) | Q16552 |
| C-X-C motif chemokine 11 (CXCL11) | O14625 | Interleukin-17C (IL-17C) | Q9P0M4 |
| Cystatin D (CST5) | P28325 | Interleukin-18 (IL-18) | Q14116 |
| Delta and Notch-like epidermal growth factor-related receptor (DNER) | Q8NFT8 | Interleukin-18 receptor 1 (IL-18R1) | Q13478 |
| Eotaxin (CCL11) | P51671 | Interleukin-20 (IL-20) | Q9NYY1 |
| Eukaryotic translation initiation factor 4E-binding protein 1 (4E-BP1) | Q13541 | Interleukin-20 receptor subunit alpha (IL-20RA) | Q9UHF4 |
| Fibroblast growth factor 21 (FGF-21) | Q9NSA1 | Interleukin-22 receptor subunit alpha-1 (IL-22 RA1) | Q8N6P7 |
| Fibroblast growth factor 23 (FGF-23) | Q9GZV9 | Interleukin-24 (IL-24) | Q13007 |
| Fibroblast growth factor 5 (FGF-5) | P12034 | Interleukin-33 (IL-33) | O95760 |
| Fibroblast growth factor 19 (FGF-19) | O95750 | Latency-associated peptide transforming growth factor beta-1 (LAP TGF-beta-1) | P01137 |
| Fms-related tyrosine kinase 3 ligand (Flt3L) | P49771 | Leukemia inhibitory factor (LIF) | P15018 |
| Leukemia inhibitory factor receptor (LIF-R) | P42702 | STAM-binding protein (STAMBP) | O95630 |
| Macrophage colony-stimulating factor 1 (CSF-1) | P09603 | Stem cell factor (SCF) | P21583 |
| Matrix metalloproteinase-1 (MMP-1) | P03956 | Sulfotransferase 1A1 (ST1A1) | P50225 |
| Matrix metalloproteinase-10 (MMP-10) | P09238 | T cell surface glycoprotein CD6 isoform (CD6) | P30203 |

| | | | |
|---|---|---|---|
| Monocyte chemotactic protein 1 (MCP-1) | P13500 | T-cell surface glycoprotein CD5 (CD5) | P06127 |
| Monocyte chemotactic protein 2 (MCP-2) | P80075 | T-cell surface glycoprotein CD8 alpha chain (CD8A) | P01732 |
| Monocyte chemotactic protein 3 (MCP-3) | P80098 | Thymic stromal lymphopoietin (TSLP) | Q969D9 |
| Monocyte chemotactic protein 4 (MCP-4) | Q99616 | TNF-beta (TNFB) | P01374 |
| Natural killer cell receptor 2B4 (CD244) | Q9BZW8 | TNF-related activation-induced cytokine (TRANCE) | O14788 |
| Neurotrophin-3 (NT-3) | P20783 | TNF-related apoptosis-inducing ligand (TRAIL) | P50591 |
| Neurturin (NRTN) | Q99748 | Transforming growth factor alpha (TGF-alpha) | P01135 |
| Oncostatin-M (OSM) | P13725 | Tumor necrosis factor (Ligand) superfamily, member 12 (TWEAK) | O43508 |
| Osteoprotegerin (OPG) | O00300 | Tumor necrosis factor (TNF) | P01375 |
| Programmed cell death 1 ligand 1 (PD-L1) | Q9NZQ7 | Tumor necrosis factor ligand superfamily member 14 (TNFSF14) | O43557 |
| Protein S100-A12 (EN-RAGE) | P80511 | Tumor necrosis factor receptor superfamily member 9 (TNFRSF9) | Q07011 |
| Signaling lymphocytic activation molecule (SLAMF1) | Q13291 | Urokinase-type plasminogen activator (uPA) | P00749 |
| SIR2-like protein 2 (SIRT2) | Q8IXJ6 | Vascular endothelial growth factor A (VEGF-A) | P15692 |

**Supplemental Table 5. Membrane and intracellular staining panel for flow cytometry.**

| Marker | Clone | Fluorochrome | Company | Dilution |
|---|---|---|---|---|
| **Surface staining** | | | | |
| CD3 | UCHT1 | V500 | BD Horizon | 1:10 |
| CD4 | HI30 | APC-Cy7 | BioLegend | 1:80 |
| CD8 | SK1 | BV650 | BioLegend | 1:40 |
| CD19 | HIB19 | BV711 | BioLegend | 1:40 |
| CD33 | WM53 | PE-Cy7 | BioLegend | 1:40 |
| CD45 | HI30 | BV605 | BioLegend | 1:10 |
| CD56 | NCAM16.2 | BV786 | BD Horizon | 1:10 |
| **Viability staining** | | | | |
| Fixable viability dye | - | eFluor™ 450 | eBiosciences | 1:1000 |
| **Intracellular staining** | | | | |
| IL-2 | MQ1-17H12 | PerCP-Cy5.5 | BioLegend | 1:20 |
| IL-6 | MQ2-13AS | FITC | Fluidigm | 1:7 |
| TNF | MAb11 | PE | BioLegend | 1:20 |
| GM-CSF | BVD2-21C11 | APC | BioLegend | 1:10 |

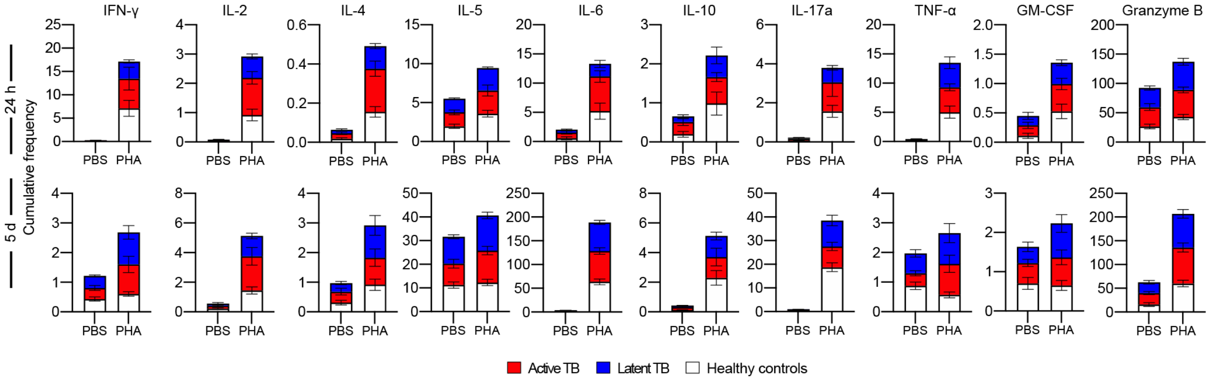


**Supplemental Figure 1. Intracellular cytokine staining after PHA stimulation.**

Frequency of cytokine-producing leukocytes with PHA stimulation after 24 hours (top) and 5 days (bottom). All donors merged (n=15/condition).

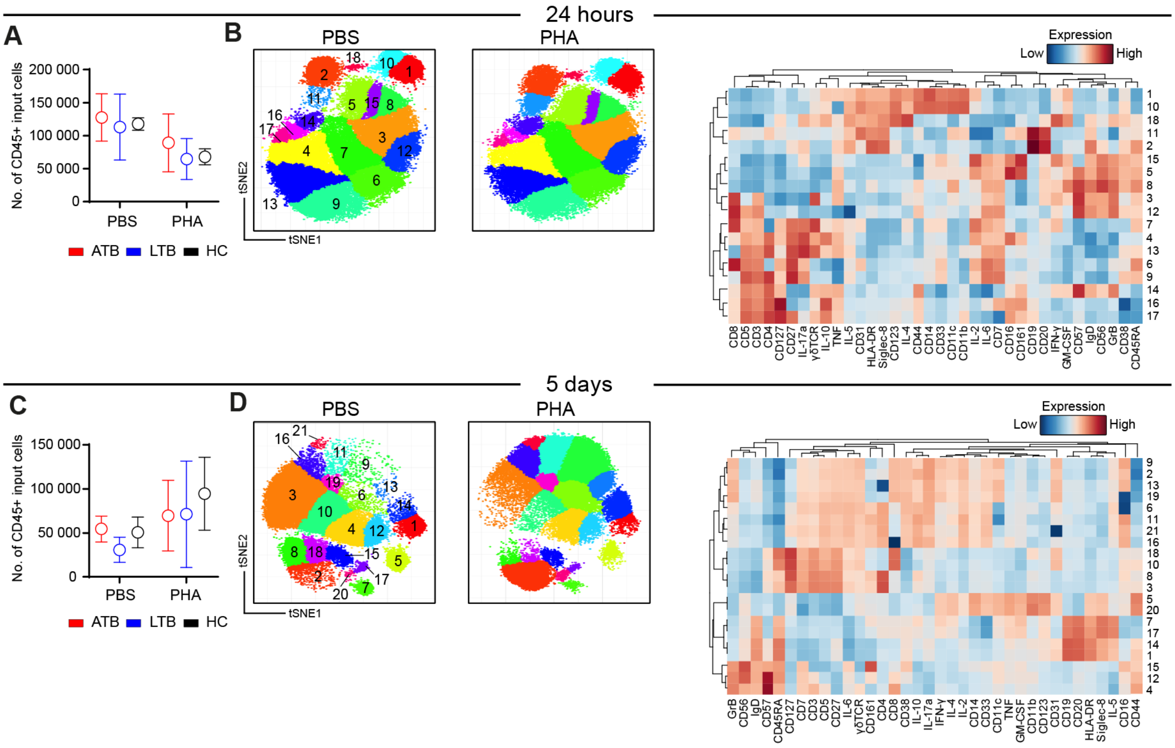


**Supplemental Figure 2. Clustering analysis of total leukocytes after 24 hours (top) and 5 days (bottom) of stimulation.**

**(A)** Numbers of CD45$^{+}$ input cells in unstimulated (PBS) and PHA-stimulated PBMCS in ATB, LTB, and HC, after 24 hours of stimulation. **(B)** Cluster identification and median heatmap expression of each marker. **(C)** Numbers of CD45$^{+}$ input cells in unstimulated (PBS) and PHA-stimulated PBMCS in ATB, LTB, and HC, after 5 days of stimulation. **(D)** Cluster identification and median heatmap expression of each marker.

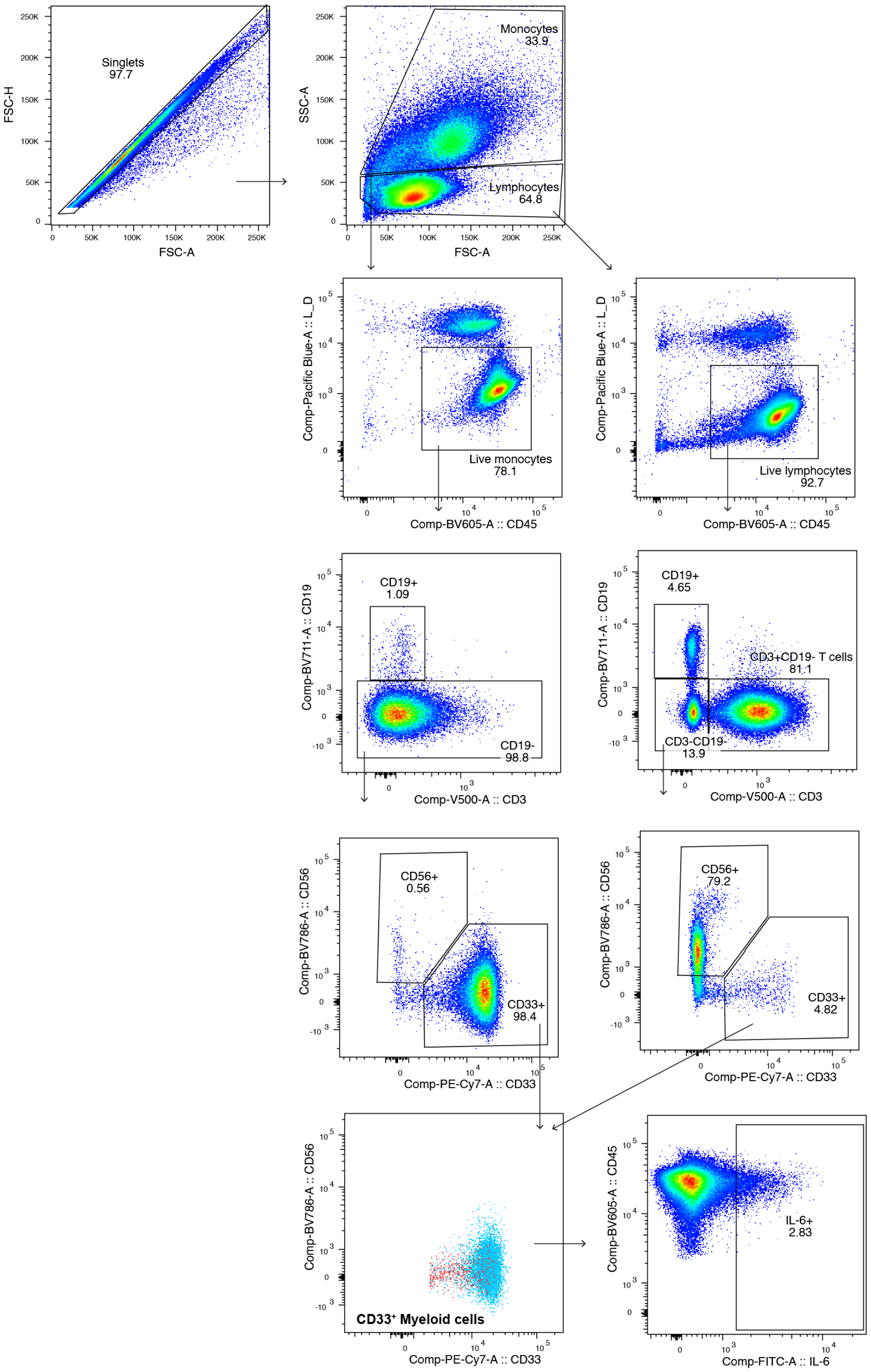


**Supplemental figure 3. Flow cytometry gating strategy for the identification of IL-6-producing-myeloid cells.** Total lymphocytes and monocytes were gated from singlets (based on FSC-H/FSC-A) based on SSC-S/FSC-A. Live lymphocytes and live monocytes were gated from total lymphocytes and total monocytes, respectively. Myeloid cells ($CD3^{-}CD33^{+}$) were gated on $CD3^{-}CD19^{-}$ both in lymphocytes and monocytes gate and merged using a Boolean gate. IL-6 producing cells were gated on myeloid cells.

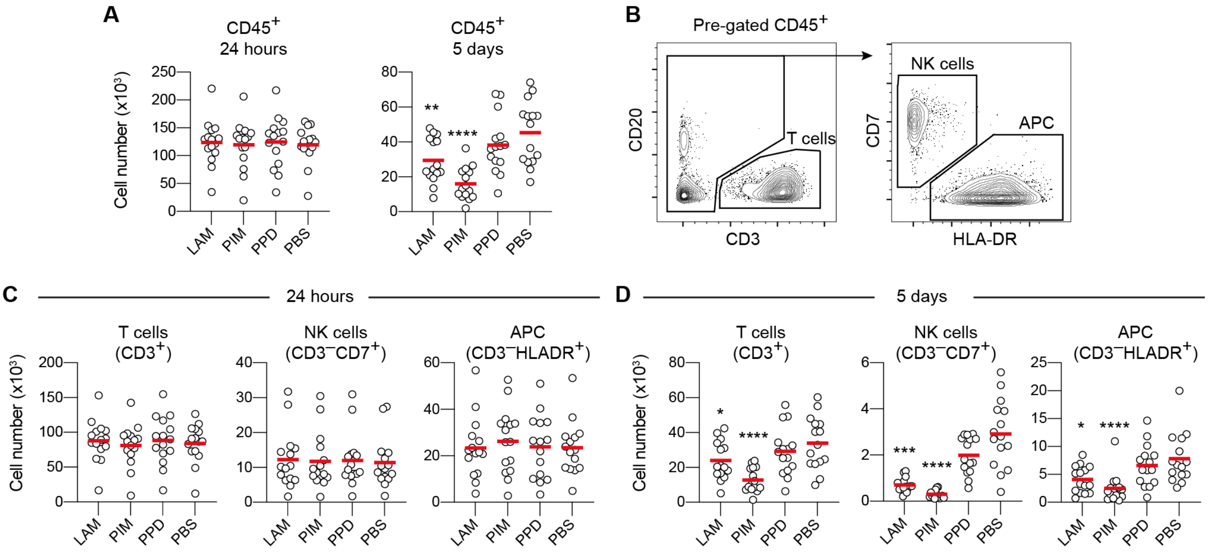


**Supplemental Figure 4. Effect of stimulation with LAM, PIM, and PPD on culture cell numbers.** (**A**) Number of total $CD45^+$ cells for each condition at 24 hours (left) and after 5 days (right) of stimulation. (**B**) Representative gating strategy to further subdivide $CD45^+$ cells into $CD3^+$ T cells, $CD3^-CD7^+$ NK cells, and $CD3^-CD7^-HLADR^+$ antigen-presenting cells (APC). (**C, D**) Number of T cells, NK cells and APCs after 24 h (**C**) and after 5 days (**D**) of stimulation. Statistics was evaluated by Friedmans test with Dunn's posttest where every group was compared with unstimulated cells (PBS). *p<0.05, **p<0.01, ***p<0.001, ****p<0.0001.

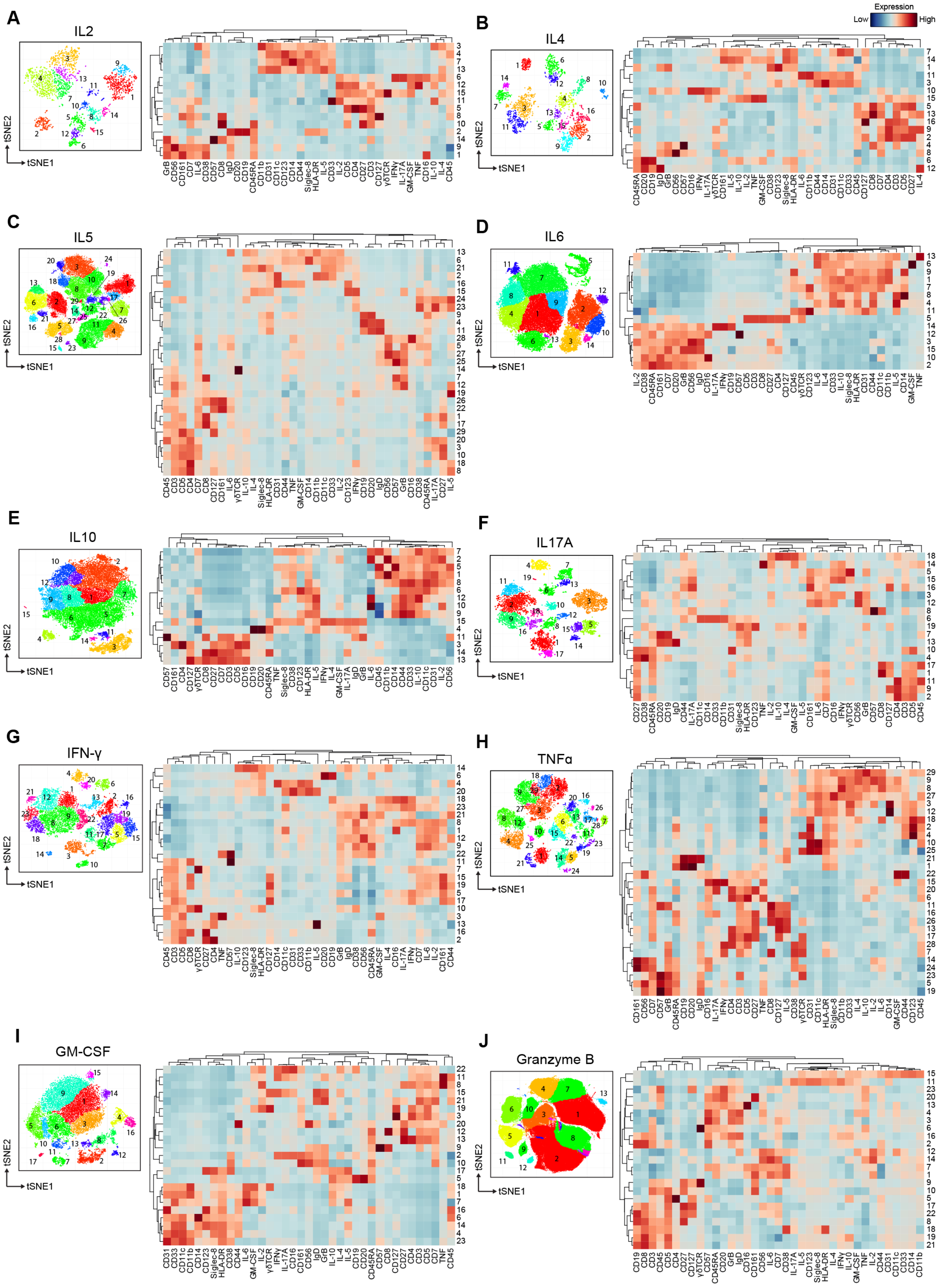

**Supplemental Figure 5. Clustering analysis of cytokine⁺ cells after 24 hours of stimulation.** Cluster identification and median heatmap expression of each marker in **(A)** IL-2, **(B)** IL-4, **(C)** IL-5, **(D)** IL-6, **(E)** IL-10, **(F)** IL-17A, **(G)** IFN-γ, **(H)** TNF-α, **(I)** GM-CSF, and **(J)** granzyme B-producing cells.

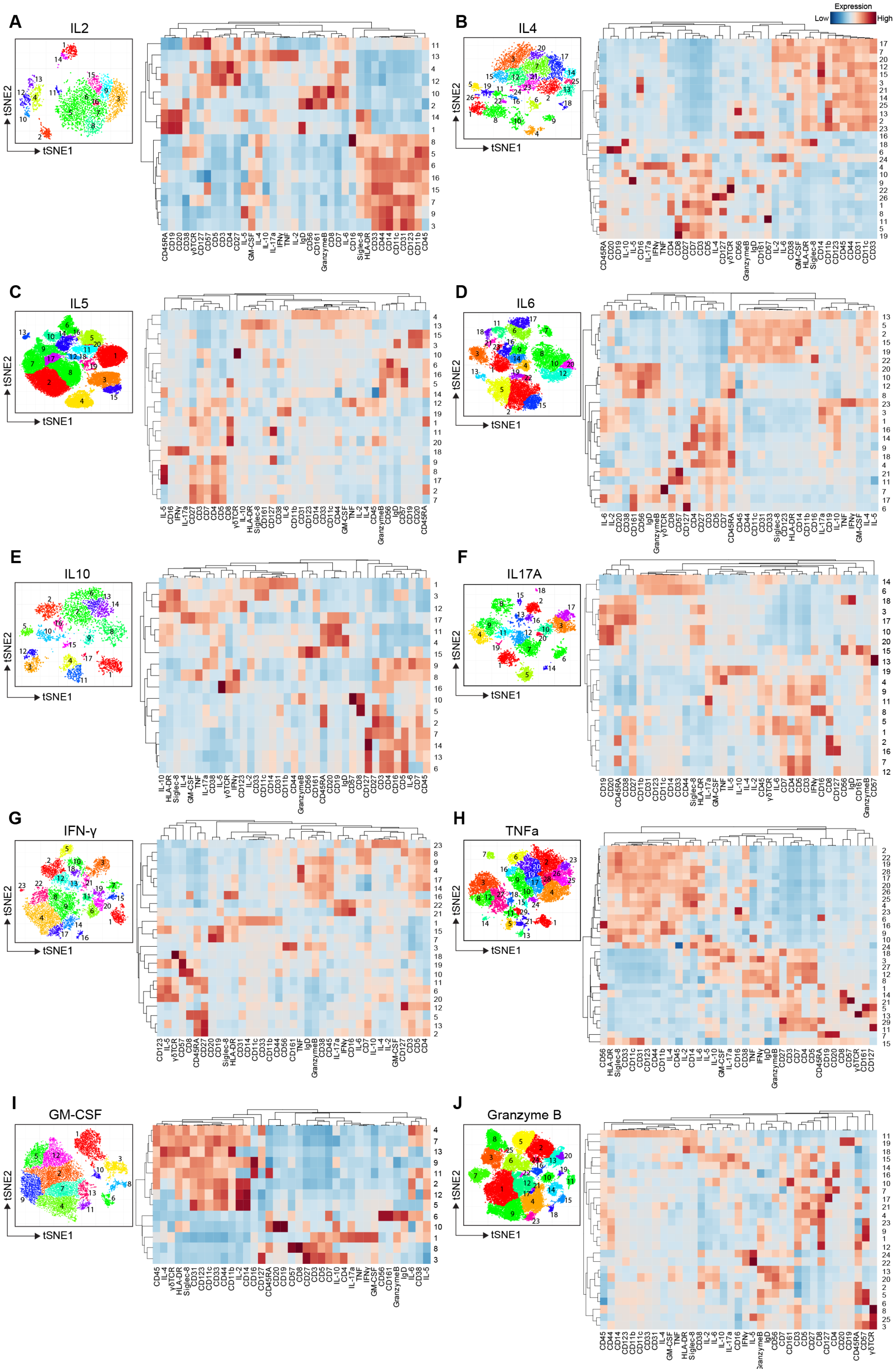
A
IL2
tSNE2
tSNE1
B
IL4
Expression
Low
High
C
IL5
D
IL6
E
IL10
F
IL17A
G
IFN-γ
H
TNFa
I
GM-CSF
J
Granzyme B

**Supplemental Figure 6. Clustering analysis of cytokine$^{+}$ cells after 5 days of stimulation.**
Cluster identification and median heatmap expression of each marker in **(A)** IL-2, **(B)** IL-4, **(C)** IL-5, **(D)** IL-6, **(E)** IL-10, **(F)** IL-17A, **(G)** IFN-γ, **(H)** TNF-α, **(I)** GM-CSF, and **(J)** granzyme B-producing cells.
